# HMGN5, an RNA or Nucleosome binding protein - potentially switching between the substrates to regulate gene expression

**DOI:** 10.1101/2022.07.15.500188

**Authors:** Ingrid Araya, Gino Nardocci, Uwe Schwartz, Sabrina Babl, Miriam Barros, Ivo Carrasco-Wong, Axel Imhof, Martín Montecino, Gernot Längst

## Abstract

The packaging of DNA into chromatin and its compaction within cells renders the underlying DNA template un-accessible for processes like transcription, replication and repair. Active mechanisms as chromatin modifying activities or the association with non-coding RNAs can de-condense chromatin, rendering it accessible for the DNA dependent processes. High mobility group proteins (HMG) are small architectural chromatin proteins that were shown to contribute to the regulation of chromatin accessibility and condensation. Here we show that HMGN5, a member of the HMGN family that is capable to de-compact chromatin exhibits a novel RNA binding domain that overlaps with its nucleosome binding domain (NBD). HMGN5 binds exclusively to nucleosomes or RNA, suggesting that molecular function relies on switching between these two substrates. We show the specific binding of HMGN5 to regulatory regions and at the same time to bind the RNA of the genes it tends to activate. Furthermore, HMGN5 co-localizes and directly interacts with CTCF, suggesting a cooperative role of both proteins in organizing higher order structures of chromatin and active chromatin domains.

## Introduction

In eukaryotic cells, DNA is highly compacted by its structural organization into chromatin and its folding into higher order structures. DNA is associated with histones, non-histone proteins and RNA, forming a dense nucleoprotein complex. Dynamic changes of the chromatin structure must occur, to enable sequence specific binding of regulatory factors and thereby regulating DNA dependent processes like transcription, replication or DNA repair ^1^. The best-characterized group of chromosomal non-histone proteins are the high-mobility-group (HMG) proteins that are present in all higher eukaryotes. Per definition, these proteins can be extracted from chromatin with 0.35 M NaCl, are soluble in 5% perchloric or trichloric acid, have a high content of charged amino acid and exhibit a molecular mass below 30 kDa ^2^. These proteins are architectural DNA binders that modulate chromatin accessibility and belong to the HMG-A, HMG box (HMGB) and HMGN families of proteins. The HMGN protein family (High Mobility Group Nucleosome Binding Domain), is only present in vertebrates and was shown to play a key role in the regulation of higher order structures of chromatin ^3^. These small proteins are tightly associated with chromatin and were shown to de-compact chromatin by interacting with nucleosomes through their highly conserved “Nucleosome Binding Domain” (NBD) ^4^. The HMGN family consists of five members, HMGN1 to HMGN5 ^4^, sharing a conserved N-terminal domain containing the Nuclear Localization Signal (NLS) and the NBD. The NBD harbors a conserved octapeptide sequence (RSARLSA) required for nucleosome binding, and HMGN proteins were shown to compete with the binding of histone H1 ^3^. The linker histone H1 binds close to the DNA entry/exit site of nucleosomes, leading to a compaction of the chromatin fiber and a general repression of gene expression ^5,6^. Thus, HMGN binding would release H1 and result in the de-compaction of higher-order structures of chromatin ^3^.

In contrast to the other HMGN members, murine HMGN5 is larger and has an a 300 aa long and highly acidic C-terminal tail (in humans - 200 aa) ^7-9^. This domain was shown to be required for tethering the mouse protein to euchromatic domains and in humans to both, euchromatic and heterochromatic domains ^9,10^. Of all HMGN members, HMGN5 has the most pronounced effect on transcriptional regulation, as deregulation of HMGN5 alters the expression of a large set of genes in cell culture and mouse models ^9-12^.

HMGN5 plays an important role in embryonic development, as HMGN5 dependent changes in chromatin structure directly affect the regulation of developmental genes ^13^. Furthermore, it was demonstrated that HMGN5 dependent global chromatin de-compaction decreases the sturdiness, elasticity and rigidity of the nucleus by a global loss of heterochromatin ^14^. Despite the functional relevance of this protein, the molecular mechanisms by which HMGN5 is targeted to its site of activity and how it regulates the changes of chromatin structure is still not known.

Chromatin associated RNAs are known to play a major role in the regulation of many chromatin-mediated processes including gene regulation, splicing and nuclear architecture ^15-18^. In human, mouse and drosophila cell lines it was also shown that chromatin associated RNAs maintain an open chromatin state, with snoRNAs and proteins like the drosophila decondensation factor DF31 playing essential roles in maintaining the accessible higher order structure of chromatin ^17,18^. A potential interplay between the HMGN proteins and chromatin associated RNA to maintain open chromatin was not addressed so far.

Here we study the functional role of HMGN5 in regulating chromatin structure and gene expression by biochemical and genome-wide methods. We identified a novel and specific RNA binding domain within HMGN5 that overlaps with its nucleosome binding domain (NBD). Binding studies revealed that HMGN5 exclusively binds to nucleosomes or to RNA, but not to both molecules at the same time. Furthermore, we showed that the RNA binding activity is a key feature of all HMGN members, highlighting potential new functions for those proteins.

The overexpression and knockdown of HMGN5 in human cell lines affected the expression of about 3000 genes respectively, with 1287 overlapping target genes. HMGN5 ChIP-seq analysis revealed the association with active regulatory regions of the genome, and in addition, HMGN5 is associated with RNA polymerase II binding sites. De-regulation of HMGN5 is strongly correlated (34%) with the transcriptional change of the asscociated genes, suggesting a direct link. Moreover, we show that HMGN5 regulated genes belong to a network of genes involved in RNA metabolic processes. CLIP-seq experiments revealed that HMGN5 binds to the nascent transcripts in proximity to its chromosomal binding sites, suggesting a dual role for HMGN5 in nuclear architecture and gene regulation. Its dual binding mode enables HMGN5 binding either to chromatin or to RNA. We found that of the 2926 identified HMGN5-bound RNAs, 622 showed HMGN5-dependent transcriptional changes in the RNA-seq after HMGN5 overexpression and 543 RNAs overlapped with genomic HMGN5 binding sites located at promoters, indicating that they are direct targets of HMGN5.

Interestingly, HMGN5 binding sites do co-localizes with CTCF binding sites *in vivo* and we show by *in vivo* and *in vitro* binding assays that HMGN5 and CTCF do interact, which suggests a cooperative role of both proteins in the organization of higher order structures of chromatin.

## Materials and methods

### Constructs and Recombinant protein purification

Full-length human HMGN5 (NM_030763.2) and deletion mutants Δ N19, Δ N20, Δ N24 and NBD were created as N-terminal GST tagged proteins by inserting the coding sequences in a modified pGEX-4T3 vector (pGEX-TEV), using KpnI and XhoI restriction sites. Mutant Δ C27 was cloned as a N-GST tagged protein using the vector pGEX-4T3 with the cloning sites BamHI and SalI. Phosphomimetic mutants S20E, S24E and S20,24E were created by site-directed mutagenesis (SDM) using the plasmid pGEX_TEV_HMGN5 as template with partially overlapping primers (all primers listed in Supplementary table 8), according to the protocol described by Zheng et al. ^19^. Mutagenesis was performed with the Q5 HF polymerase (NEB) with 16 cycles of amplification and 1 min extension per kbp at 72°C. After PCR, the plasmid was purified with the QIAquick PCR Purification Kit (Qiagen) and treated with 10U of DpnI at 37°C for 1h to degrade the methylated parental plasmid. After incubation, DpnI was heat-inactivated at 80°C for 20 min. All constructs were verified by sequencing before bacterial transformation.

For LacI/LacO tethering of HMGN5, the plasmid pSV2_GFP-LacI was obtained from Dr. Karsten Rippe. The psV2_HMGN5-GFP-LacI construct was created by inserting HMGN5 CDS in a modified version of pSV2_GFP-LacI (gblock_pSV2_GFP_LacI) using EcoRI and XhoI restriction sites.

Proteins were expressed in BL21 *E*.*coli* competent cells, by induction with 1mM IPTG at an OD_600_ of 0.6. Proteins were purified with 4% Glutathione Agarose (Jena Biosciences) according to the manufacturer’s instructions. Proteins were dissolved in interaction buffer (20 mM Tris-HCl pH 7.6, 1.5 mM MgCl_2_, 0.5 mM EGTA, 10% glycerol, 100 mM KCl) and quantified using the Qubit fluorometer (Invitrogen). All plasmids used in this study are listed in Supplementary table 9.

### Nucleosome assembly

Mono-nucleosomes were reconstituted by the salt gradient dialysis method on Cy3 labeled PCR fragments containing the 601 nucleosome positioning sequence ^20,21^. Histones were purified from chicken blood. Standard assembly reactions (50µl volume) were performed at histone:DNA ratios of 1.4:1 and 2:1 (Supplementary Figure 11), with 1µg of the purified, fluorescently labeled DNA and 200 ng/µl BSA in High salt buffer (10 mM Tris, pH 7.6, 2 M NaCl, 1 mM EDTA, 0.05% NP-40, 2 mM ß-mercaptoethanol). The assembly took place for 16-20h at room temperature by decreasing continuously the salt concentration to 200 mM NaCl. *In vitro* assembled mono-nucleosomes were analyzed by native PAGE (6% in 0.4x TBE) and visualized on a fluorescence imager (Typhoon).

### *In vitro* interaction and competitive EMSA

Binding assays were performed by electrophoretic mobility shift assay (EMSA) and Microscale thermophoresis (MST, NanoTemper Munich). Dilution series (1.5 fold) of the HMGN5 proteins were prepared, starting from 2 µM protein in reaction buffer (100 mM KCl, 20 mM Tris-HCl pH 7.6, 1.5 mM MgCl_2_, 0.5 mM EGTA, 10% glycerol, 0.05% IGEPAL CA-630).Each protein sample was mixed with 50 nM of fluorescently labeled nucleic acids (Supplementary table 10) or 50 ng/µl of fluorescently labeled mono-nucleosomes in reaction buffer. The samples were incubated for 15 min at room temperature, and then subjected to EMSA or MST analysis (below). EMSAs were loaded on 6 to 8% native PAA gels and run in 0.4x TBE for 30 min at 100V (protein-nucleic acids interactions), or for 1h at 100V when analyzing protein-nucleosome interactions. Gels were documented using a fluorescence imager (Typhoon 9500, GE Healthcare Life Sciences, Munich, Germany). Nucleosome-binding activity test of the recombinant HMGN5 is shown in Supplementary Figure 12.

To perform competitive binding assays between RNA and nucleosome, protein-nucleosome complexes were kept at constant concentration (2 μM protein and 50 ng/µl of nucleosome) and increasing amounts of non-labeled fragmented cellular RNA was added (300ng, 700ng, 1μg and 2μg). The reactions were loaded onto 6% native PAA gel and run 1h at 100V. Gels were documented using a Typhoon 9500 device.

### MicroScale Thermophoresis (MST) analysis

MicroScale Thermophoresis (MST) binding experiments were carried out essentially as previously described ^22^. MST measurements were performed in reaction buffer with a range of protein concentrations at 40% MST power, 20% LED power in standard capillaries at 25°C on a Monolith NT.115 device (NanoTemper Technologies). The data was analyzed using the MO.affinity analysis software (V2.3, NanoTemper Technologies) and binding reactions were determined by analysing changes in Temperature Related Intensity Changes (TRIC-effect). Data was quantified and fitted using the Hill model, as the binding curves revealed cooperative effects. To calculate the fraction bound, the ΔF_norm_ value of each point is divided by the amplitude of the fitted curve, resulting in values from 0 to 1 (0 = unbound, 1 = bound), and processed using the Sigmaplot 12.0 software.

### Western blot and immunofluorescence microscopy

20µg of protein extracts or 10% Co-IP samples were resolved on 10-12% SDS–PAGE and proteins were transferred to PVDF membranes using a semi dry Western blot system (Bio-Rad Trans-Blot SD). Membranes were blocked in 5% non-fat milk in PBS-T (1x PBS, 0.1% Tween 20) for 1h at RT. Incubation with primary and secondary antibody was performed in blocking solution. The primary antibody was incubated for 1h at room temperature with 3 washing steps of 3-5 min in PBS-T, prior to addition of the secondary antibody. The SuperSignal WEST Dura Kit (Pierce) was used for chemiluminescent detection and signals were recorded with a LAS-3000 Imager (Fujifilm).

For immunofluorescence analysis, cells were seeded on coverslips in 24-well plates. Cells were fixed with 4% PFA in PBS for 10 min at room temperature. Fixed samples where then washed 3 times for 3 min with 0.01% Triton X-100 in PBS, one time with 0.5% Triton X-100 in PBS and twice with PBS. Cells were incubated with primary and secondary antibodies (in 4% Albumin Bovine Fraction V (Serva), PBS-T) for 45 min and 30 min, respectively. Three washing steps with 0.1% PBS-Tween were performed after incubation of the primary antibody. DNA was counterstained with 50 ng/ml DAPI in PBS-T for 5 min. Images were acquired with an Axiovert 200M microscope or with confocal microscope Leica SP8.

The following primary antibodies were used for Western blot: HMGN5 antibody (HPA000511, Sigma) at a dilution of 1:2500, β-actin antibody (A2066, Sigma) at a 1:5000 dilution, CTCF (sc-28198, Santa Cruz) at a 1:1000 dilution. Commercial standard Anti-rabbit HRP conjugated secondary antibody was used at 1:5000 dilution. For immunostaining, the anti HMGN5 antibody HPA000511 was used at a dilution of 1:500. The secondary antibody anti-Rabbit-DL488 (Jackson Laboratory) was used at 1:200 dilution. Double staining for confocal imaging was performed with the CTCF antibody sc-271474 (Santa Cruz) and HMGN5 HPA000511 (Sigma) both at 1:300 dilution. The respective secondary antibodies anti-Rabbit Rhodamine red-x 650-580 and anti-Mouse Alexa 488 were used at 1:500 dilution.

### Microscopical analysis of protein co-localization in cells

Immunofluorescence acquisition was performed on a Leica TCS SP8 (Leica Microsystems Inc., Mannheim, Germany) laser scanning confocal equipped with HC PL APO CS2 NA 1.4 63X oil optical magnification. Pinhole size was set to 1 AU. Samples were illuminated with 405, 488 and 552 nm LED lasers. 1024×1024 pixel (xy 36,1 nm/pixel) images were acquired using 3 PMT detectors. Emitted fluorescence was split with dichroic mirrors DD488/552. Hoescht (Sigma-Aldrich, ST. Lois, MO, USA) acquisition parameters were 410nm - 483nm (emission wavelength range), 750 (gain) -1 (offset). Alexa Fluor 488 (InvitrogenTM, ThermoFisher Scientific,Waltham, MA, USA) acquisition parameters were: 505-550nm (emission wavelength range), 850 (gain) -1 (offset). Rhodamine acquisition parameters were: 587-726 nm (emission wavelength range), 850 (gain) and -1 (offset). For z-stack imaging, a total of 50 slices were acquired with a 100nm z-step size.

All images were deconvolved with Huygens Professional version 18.02 (Scientific Volume Imaging, The Netherlands, http://svi.nl), using the CMLE algorithm, with SNR:15 and 40 iterations. The Co-localization Analyzer (Scientific Volume Imaging, The Netherlands, http://svi.nl), was used to perform co-localization studies.

Co-localization degrees were assigned as described in Zinchuk et al. ^23^. The set includes Pearson’s description for five variables: -1 ∼ -0.27 “Very weak”, -0.26 ∼ -0.09 “Weak”, 0.1 ∼ 0.48 “Moderate”, 0.49 ∼ 0.84 “Strong”, and 0.85 ∼ 1.0 “Very strong”, which was used as a standard to describe the results of quantitative co-localization studies ^23^. Controls were generated by flipping HMGN5 channel 180 degrees to measure incidental co-localization.

### Establishement and maintenance of stable cell lines

The stable cell line expressing HMGN5 (HMGN5-FlpIn) was created using the T-REx™-293 Flp-In system (Life Technologies) according to the manufacturer’s protocol. The coding sequence of HMGN5 (NM_030763.2) was inserted in the expression vector pcDNA5/FRT/TO_GFP, using the HindIII and XmaI restriction sites, to obtain a C-terminal GFP fusion construct. T-REx™-293 Flp-In cells were transfected with the pOG44 and pcDNA5/FRT/TO© constructs in a 9:1 ratio using FuGENE® HD Transfection Reagent. After a 24h incubation at 37°C, cells were trypsinized and expanded in 10cm plates in DMEM/10% Tetracycline-free FBS/ 100 μg/ml Hygromycin/ 10 μg/ml Blasticidin. After 10-12 days incubation, Hygromycin-resistant foci were expanded and tested for protein expression using 1 μg/ml Doxycycline. Clones exhibiting homogeneous protein expression in immunofluorescence were pooled and used for the experiments.

For maintenance, HMGN5-FlpIn cells and a control cell line expressing GFP (GFP-FlpIn) were grown in low glucose DMEM GlutaMAX™ supplemented with 10% Tetracycline-Free FBS (Biochrom), 100 µg/ml Hygromycin and 10 μg/ml Blasticidin, at 37°C and 5% CO_2_. Protein induction was performed with 1 μg/ml Doxycycline in DMEM/ 10% Tetracycline-Free FBS.

### Chromatin de-compaction analysis

U2OS cells containing a stable integration of the LacO array in a highly compacted telomeric region ^24^ (obtained from Karsten Rippe’ laboratory) were grown in low glucose DMEM GlutaMAX™ supplemented with 10% FBS (Gibco), at 37°C and 5% CO_2_. Cells were transfected with the construct pSV2_GFP_LacI_HMGN5 or with the control plasmid pSV2_GFP_LacI for transient expression of HMGN5_GFP_LacI or GFP_LacI respectively, to analyze the tethering to the LacO array. 24h post transfection the cells were fixed with 4% PFA and the DNA was counterstained with DAPI. Cells were visualized by confocal microscopy using a Leica SP8 device.

### Knockdown of HMGN5 by RNA interference

HMGN5-FlpIn cells were cultured in low glucose DMEM medium without antibiotics. At 70% confluence, cells were transfected with a final concentration of 40 nM of a SmartPool siRNA against HMGN5 or a non-targeting SmartPool siRNA (purchased from Dharmacon) using Lipofectamine RNAiMax (Invitrogen) according to manufacturer’s instructions. 24 hours after transfection the cells were harvested and used for RNA sequencing. HMGN5 protein levels were analyzed by Western blot.

### cDNA synthesis and quantitative PCR

Cellular RNA for cDNA synthesis was purified with the NucleoSpin RNA Kit (MACHEREY-NAGEL) according to manufacturer’s instructions. cDNA was prepared from 500ng total RNA using the SuperScript II reverse transcriptase (Invitrogen) and random hexamers (Invitrogen) following the manufacturer’s instructions. RT-qPCR validation of the RNA-seq data was performed using oligonucleotide primers for the candidate mRNAs ZCCHC12, VGF, RPL30 and EGR1. GAPDH was used as reference gene for quantification (list of primers in supplementary table 8). Three independent biological replicates were used for each, HMGN5 overexpression, knock-down and control cells. qPCR reactions were performed with the SYBR® Green qPCR Kit (Qiagen) as described ^25^. To define statistical differences between mRNA levels from knock-down, overexpression and control groups Two-way ANOVA and Dunett’
ss post-test were used ^26^.

### RNA-seq library preparation and data analysis

Total RNA was isolated in biological duplicates from the HMGN5-FlpIn cell line after 24h induction, 24h knock-down and from non-treated control cells (70% confluent 10 cm culture dish per replicate). RNA was purified with the NucleoSpin RNA Kit (MACHEREY-NAGEL) according to the manufacturer’s instruction. Remaining genomic DNA was eliminated with TURBO DNA-free™ DNase (Ambion). At least 3µg RNA per condition, determined by Qubit RNA HS Assay Kit, were used for library preparation and high throughput sequencing by the EMBL GeneCore Facility in Heidelberg (Dr. Vladimir Benes). The sequencing was performed on the Illumina Hiseq2000 platform with a read length of 50bp paired-end. Quality control was carried out with the FastQC package. All pre-processing steps were performed using the SAMtools or BEDtools packages. Transcripts were mapped to the hg19 reference genome using the STAR aligner using standard settings. Differential gene expression analysis of HMGN5 over-expression and knock-down over the non-treated control was performed with the DEseq2 package from R (v3.1.2) ^27^.

### ChIP-seq experiment and data analysis

Chromatin immunoprecipitation (ChIP) was performed of non-induced HMGN5-FlpIn control cells, HMGN5-FlpIn cells induced for 24h (two biological replicates each) and GFP-FlpIn cells (with or without induction with doxycycline for 24h). Cells were grown to 70-80% confluence in 15 cm culture plates and crosslinked with 0.8% formaldehyde in 1x PBS for 10 min at RT. The reaction was quenched by the addition of 125 mM glycine (final concentration), and subsequently washed with ice-cold PBS. Cells were lysed in 900 μl SDS lysis buffer (1% SDS, 50 mM Tris-HCl pH 8.0, 20 mM EDTA, protease inhibitors) for 10 min on ice. Chromatin was fragmented with a Bioruptor (Diagenode) to obtain a DNA fragment length between 200 and 500 bp. The samples containing the chromatin were diluted with ChIP Low salt buffer (20 mM Tris-HCl pH8.0, 150 mM NaCl, 2 mM EDTA, 1% Triton X-100, Protease inhibitors) and incubated on a rotating wheel with 10 µl of precleared GFP-Trap Agarose beads (Chromotek) for 2 hours at 4°C. Beads were washed twice with ChIP Low salt buffer, once with ChIP High salt buffer (20 mM Tris-HCl pH 8.0, 500 mM NaCl, 2 mM EDTA, 1% Triton X-100), once with ChIP LiCl buffer (10 mM Tris-HCl pH 8.0, 250 mM LiCl, 1 mM EDTA, 1 % IGEPAL CA-630, 1 % (w/v) sodium deoxycholate) and once with 500 µl EB buffer (Qiagen). Beads were treated with 300 µg/ml RNase A, followed by Proteinase K treatment (100 µg/ml, 1h at 55°C) and de-crosslinking overnight at 65°C. The DNA was purified and quantified with the Qubit® dsDNA BR Assay Kit (Invitrogen). DNA samples were used to prepare sequencing libraries using the NEBNext® ChIP-seq Library kit following the manufacturer’s recommendations. Deep sequencing was performed on an Illumina HiSeq 1000 platform, using 50bp single-end parameter, at the KFB (Kompetenzzentrum für fluoreszente Bioanalytik) in Regensburg. The quality of the sequencing reaction was analyzed with the FastQC software. HMGN5 non-induced samples were used as background for each individual HMGN5 induced sample. The sequences were aligned to the hg38 reference genome using Bowtie2 with standard settings ^28^. PCR duplicates were removed with Picard (http://broadinstitute.github.io/picard/). The reads from each group of samples were merged to call peaks using MACS2 software ^29^. Peak calling resulted in 8952 peaks. Peak finding was performed applying broad peak calling parameters. Gene ontology enrichment analysis was performed using the R package GOstats ^30^ and ClueGo plug-in (v2.3.3) ^31^. De novo Motif finding was performed using the ‘findMotifsGenome.pl’ command from HOMER (v4.9) ^32^.

Analysis of histone modifications at HMGN5 peaks was conducted using the computeMatrix and plotHeatmap functions from the deeptools package.^33^

### CLIP-seq experiment and data analysis

HMGN5-FlpIn cells and the control GFP-FlpIn cells were used for UV-crosslinking RNA immunoprecipitation using a modified protocol. 70-80% confluent cells were induced with 1 µg/µl Doxycycline for 6 hours. Cells were UV-crosslinked with 150 mJ/cm^2^ at 254 nm (stratalinker UV crosslinker, Stratagene) on ice, and harvested in 1.5 ml ice-cold PBS. The samples were diluted with Low salt buffer (20 mM Tris-HCl pH7.5 250 mM NaCl, 1 mM MgCl_2_, 0.025% SDS, 0.05% IGEPAL CA-630, RNAsin, protease inhibitors) and sonicated 15 min high intensity 30s ON/OFF in a Bioruptor device. Cells were pelleted and incubated with 10 µl of pre-equilibrated GFP-Trap_M beads (Chromotek) 2 hours at 4°C with rotation. After incubation, the samples were washed three times with 700 µl Low salt buffer, then twice with 700 µl High salt buffer (50 mM Tris-HCl pH 7.5, 500 mM NaCl, 1 mM MgCl2, 0.1% SDS, 0.05% NP-40, RNasin, protease inhibitors) and finally once with 500 µl of Elution buffer (10 mM Tris-Cl, pH 7.5). RNA was partially digested on beads in 500 µl of EB using RNase A/T1 (Thermo Scientific) at 37°C for 10 min and washed once with Low salt buffer and once with Elution buffer. DNA was removed with DNase Q1 for 1h at 37°C. Samples were treated with 300 µg/ml Proteinase K for 2 hours at 37°C. The RNA was purified from the beads using Trizol and resuspended in 20µl Elution buffer. Eluted RNA was analyzed using the Agilent RNA 6000 Pico Kit in a Bioanalyzer. Stranded libraries were prepared using the Ovation Universal RNA-seq kit (Nugen) according to the manufacturer’s instructions. Library size distribution was analyzed using a High Sensitivity DNA kit (Agilent) in a Bioanalyzer. High-throughput sequencing was performed on an Illumina HiSeq 1000 platform using 50bp single-end parameter.

CLIP-seq reads were aligned to the hg38 human reference genome using the Bowtie software (v2.1.1). The alignments were exported as BAM files for visualization and indexes were created using the BEDtools utilities (v2.24.0). Peak finding was performed using the ‘findPeaks’ command of Homer (v4.9), combining the alignment of all three HMGN5 samples, using as background a combined alignment of the GFP2 and GFP3 samples. The number of reads per transcript was counted using HTSeq (v0.11.2) ^34^. The enrichment was estimated using using SARTools (v1.6.9) ^35^ and DESeq2 (v1.24) ^27^ R packages. De novo Motif finding was performed using the ‘findMotifsGenome.pl’ command from Homer (v4.9). Peaks were annotated using the ‘annotatePeaks.pl’ command from Homer, using default setting and stats option.

To analyze the distribution of HMGN5 binding sites along chromatin states, the ChromHMM 18-state model of HEK293T was used (ENCODE accession ENCFF071AXS). Chromatin state annotations were assigned to the corresponding HMGN5 ChIP-/ CLIP-seq peaks. In case a HMGN5 peak overlapped with more than one chromatin state annotation, the chromatin state with the largest overlap was taken. Peaks located in regions without any reliable chromatin state annotation, were marked as *no Annotation*.

### Co-immunoprecipitation and quantitative mass spectrometry

70% confluent HMGN5-FlpIn cells and the control GFP-FlpIn cells, grown in 15 cm plates, were induced with 1 μg/ml Doxycycline for 8h (3 plates from each cell line). Cells were washed once with PBS and harvested in 2ml ice-cold PBS. The cells were lysed using 1 ml of Co-IP Lysis buffer (20 mM Tris-HCl pH 7.5, 150 mM NaCl, 3 mM CaCl_2_, 0.5% IGEPAL CA-630, Protease inhibitors) containing MNase/Benzonase or only MNase, on ice and incubated 10 min at 37°C. The reaction was stopped with 20 mM EDTA. 500 μl of lysate per biological replicate were diluted with 500 μl Co-IP Wash buffer 1 (20 mM Tris-HCl pH 7.5, 150 mM NaCl, 0.5 mM EDTA, 0.5% IGEPAL CA-630, protease inhibitors) and incubated with 15 μl of GFP_Trap magnetic beads (Chromotek) for 1h at 4°C with gentle rotation. Beads were washed 3 times with 700 μl ice-cold Co-IP Wash buffer 1, and 3 times with 100 μl ice-cold Co-IP Wash buffer 2 (50 mM Tris-HCl pH8). The samples were placed on a magnetic rack for 2-3 min in between each step to remove supernatant. After immunoprecipitation, the independent biological replicates were digested on-bead and subjected to mass spectrometry. Label-free quantitative proteomics was performed by Liquid Chromatography Tandem Mass Spectrometry (LC-MS/MS) at the Protein Analysis Unit of the Biomedical Center (Ludwig Maximilian University, Munich). The data was processed with the Perseus software to calculate Intensity-based absolute quantification (iBAQ). The UniProt database was used for protein identification. Protein enrichment in the HMGN5 samples compared to the GFP control was calculated by Log2 fold change (Log2 FC) normalization over the GFP control (log2 FC IBAQ HMGN5-GFP). The significance was evaluated using t-test and limma test, both adjusted for multiple comparisons. Proteins were considered enriched in the HMGN5 samples over GFP only when Log2 fold change was ≥2 and with statistical significance (p-value <0.05) in both, t-test and limma test.

### Gene Ontology analysis and data visualization

Gene Ontology Enrichment Analysis of ChIP-seq was performed using the GOstats (v2.50.0) R package ^30^ and ClueGo plug-in (v2.3.3) ^31^ from the Cytoscape software (v.3.5.1). The significance of the enrichments was obtained with a hypergeometric test and the p-value correction with the Bonferroni step-down method (p-value <0.005). The selected leading terms correspond to the most significant term in each group. GO analysis of clustered HMGN5 peaks was performed with GREAT software (http://great.stanford.edu/public/html/splash.php). When corresponding, plots were performed using the Sigmaplot software (v12.5). Visualization of grouped datasets was performed with the BioVenn tool ^36^.

Gene Ontology (GO) analysis of CLIP-seq data was carried out using the GOstats (v2.50.0) R package ^30^.

## Data availability

The data generated in this study has been deposited in Gene Expression Omnibus (GEO) under the accession number GSE166103 (https://www.ncbi.nlm.nih.gov/geo/query/acc.cgi?acc=GSE166103).

Cytoplasmic and Nuclear RNA data from HEK293 cells was downloaded from GEO accession number GSE68671 ^37^. HEK293 ChIP-seq and DNase-seq bigWig tracks were downloaded from ENCODE repository ^38^ (Accession: H3K27ac - ENCFF885SUR; H3K9me3 - ENCFF526FQB; H3K4me3 - ENCFF439DDQ; H3Kme1 - ENCFF274LAP; H3K36me3 - ENCFF458PUF; DNase-seq - ENCFF529BOG; POLR2A - ENCFF000WYF; CTCF - ENCFF128UTY). ChromHMM 18-state model of HEK293T cells was obtained from the ENCODE repository (accession ENCFF071AXS).

## Results

### HMGN5 is a RNA binding protein

We previously demonstrated that the protein Df31 (decondensation factor 31), specifically interacts with RNA and maintains accessible higher order structures of chromatin ^18^. In humans, the high mobility group protein HMGN5 is conserved to Df31 (30% aa identity), also localizes to the nucleus and the nucleolus, and maintains euchromatic regions accessible ^9,11^. As HMGN5 resembles the functionality of Df31 we ask whether its activity is also mediated by RNA binding. To explore the RNA binding capabilities of HMGN5 we quantified the interaction of recombinant human HMGN5 (Figure 1a, Supplementary Figure 1a) with fluorescently labeled single-stranded nucleic acid sequences by electromobility shift assays (EMSA, Figure 1b) and microscale thermophoresis (MST, Figure 1c) ^39^. The ssRNA and ssDNA molecules used in the assays have the same sequences, lengths between 29 and 38 nt and different GC contents.

**Figure 1.**
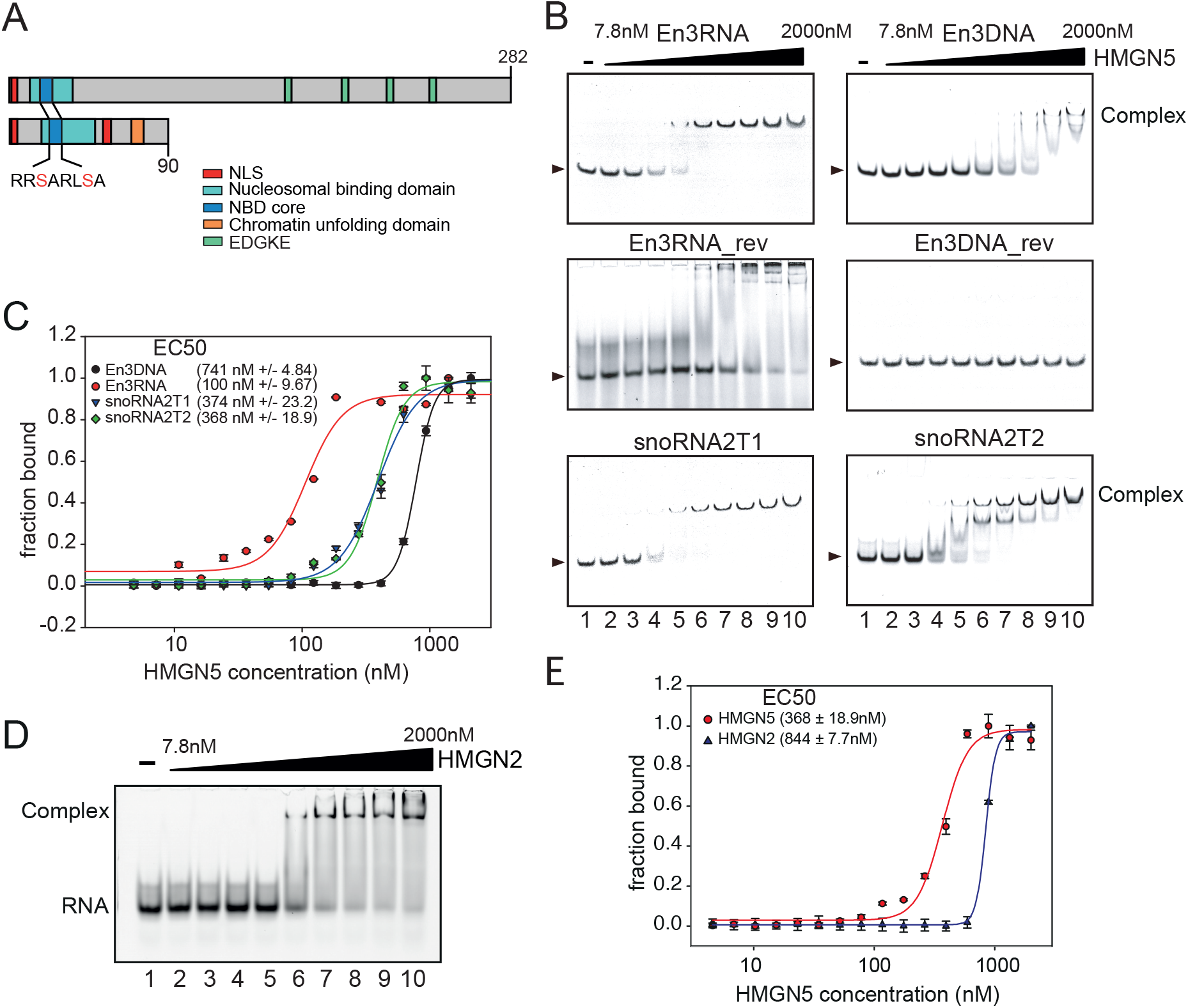
HMGN5 has a specific RNA binding ability. (A) Comparison between human HMGN5 and HMGN2. Each functional domain is represented in the scheme. The nuclear localization signal (NLS) is indicated in a red box. The nucleosomal binding domain (NBD) and the NBD core containing the conserved octapeptide RRSARLSA are represented in cyan and blue boxes respectively. Serine 20 and 24 (S20 and S24) of this sequence are highlighted in red. The orange box in HMGN2 represents the chromatin-unfolding domain, and the green boxes represent the repetitive sequence EDGKE in HMGN5. The last amino acid of HMGN5 (amino acid 282) and HMGN2 (amino acid 90) are indicated in the scheme. (B) HMGN5 binding to different single-stranded nucleic acids analyzed by EMSA experiments. Each fluorescently labeled nucleic acid was kept at a constant concentration of 50 nM. 1.5 fold titrations ranged from 0-2 μM of HMGN5 were used. Arrows indicate the position of the unbound nucleic acid molecule. (C) The protein-nucleic acid interactions were quantified by Microscale thermophoresis (MST) using the same conditions described in B. The binding curve was adjusted with the Hill equation and the EC50 value for each interaction was determined. (EC50 +/-standard deviation; n=3). The binding interaction by MST was measured for the ssRNAs En3RNA, snoRNA2T1, snoRNA2T2 and the ssDNA En3DNA. (D) Interaction of HMGN2 with the Cy5-labeled snoRNA2T2 analyzed by EMSA. 1.5 fold titrations ranged from 0-2 μM of protein were used. The RNA was used at 50 nM concentration. (E) Analysis of HMGN2-RNA interaction by MST. The binding curve was determined according to the Hill equation and the EC50 value was calculated (indicated in the nM range ± standard deviation, n=3). The binding was compared to the HMGN5-snoRNA2T2 interaction (from Figure 1C).

HMGN5 exhibits a specific, high affinity RNA binding activity giving rise to defined nucleoprotein complexes in EMSA (Figure 1b). The interaction of HMGN5 with snoRNA2T2 produces two defined bands, suggesting that the slower migrating complex is bound by at least two molecules of HMGN5 that bind specific and simultaneously to the RNA.

The specific binding of HMGN5 to RNA is a novel function of the protein, and it binds significantly better to RNA than to ssDNA of the same sequence. Binding affinities were quantified by MST, revealing a 7 times higher binding affinity of HMGN5 for certain RNA over ssDNA sequences, whereas other ssDNA sequences were not bound at all (Figure 1b, c). The protein exhibited cooperative binding curves with an affinity of 100 nM (EC50) for the RNA sequence En3 and an affinity of 741 nM for the respective ssDNA sequence. The differences in binding affinity were even more striking when comparing the reverse sequences of En3 (En3RNA_rev and En3DNA_rev, Figure 1b). HMGN5 showed clear binding of the RNA molecule, but failed to bind to the ssDNA in EMSA. Here, we describe the discovery of a novel RNA binding activity for HMGN5, which is accompanied by a significantly lower ssDNA binding affinity.

To test whether RNA binding is a general feature of HMGN family members, we tested the interaction of the human recombinant HMGN2 protein (Figure 1 and Supplementary Figure 1b) with RNA (Figure 1d and 1e). The results clearly showed the formation of specific RNA-HMGN2 complexes with intermediate affinities (EC50 844±7.7 nM), as revealed by EMSA and MST, respectively (Figure 1d and 1e). Additionally, we tested the binding of HMGB1 (Supplementary Figure 1c) to RNA (Supplementary Figure 1d), also a member of the HMG super family that is able to interact with chromatin and to regulate nuclear processes like transcription, replication or DNA repair ^40^. In contrast to the HMGN proteins, HMGB1 failed to interact with RNA (Supplementary Figure 1d), indicating that the RNA binding activity is an intrinsic feature of the HMGN family. These findings suggest a potential role of RNA molecules in the HMGN mediated de-condensation of chromatin.

### The nucleosome binding domain is required but not sufficient for RNA binding

Members of the HMGN family possess the Nucleosome Binding Domain (NBD), which is essential for nucleosome interaction. It was shown that two critical serine residues in the NBD are required for nucleosome binding (S20 and S24 in human HMGN5), which are phosphorylated during metaphase, releasing the protein from chromatin ^9,10,41^.

We performed a mutational analysis to identify the RNA binding domain and show that the NBD contributes to RNA binding (Figure 2). We analyzed the RNA binding capabilities of GST-tagged deletion mutants and three phosphomimetic mutants targeting the S20 and S24 residues in the NBD of HMGN5 (Figure 2a, b). EMSA experiments with the RNA snoRNA2T2 showed that the deletion of the first 19 amino acids (Δ N19) markedly reduced the affinity of the protein for RNA, and deletion of the first 24 amino acids (Δ N24) including serines S20 and S24, completely abolished RNA binding (Figure 2c). Unexpectedly, the NBD region seems to be required for RNA binding as it is for nucleosome binding.

**Figure 2.**
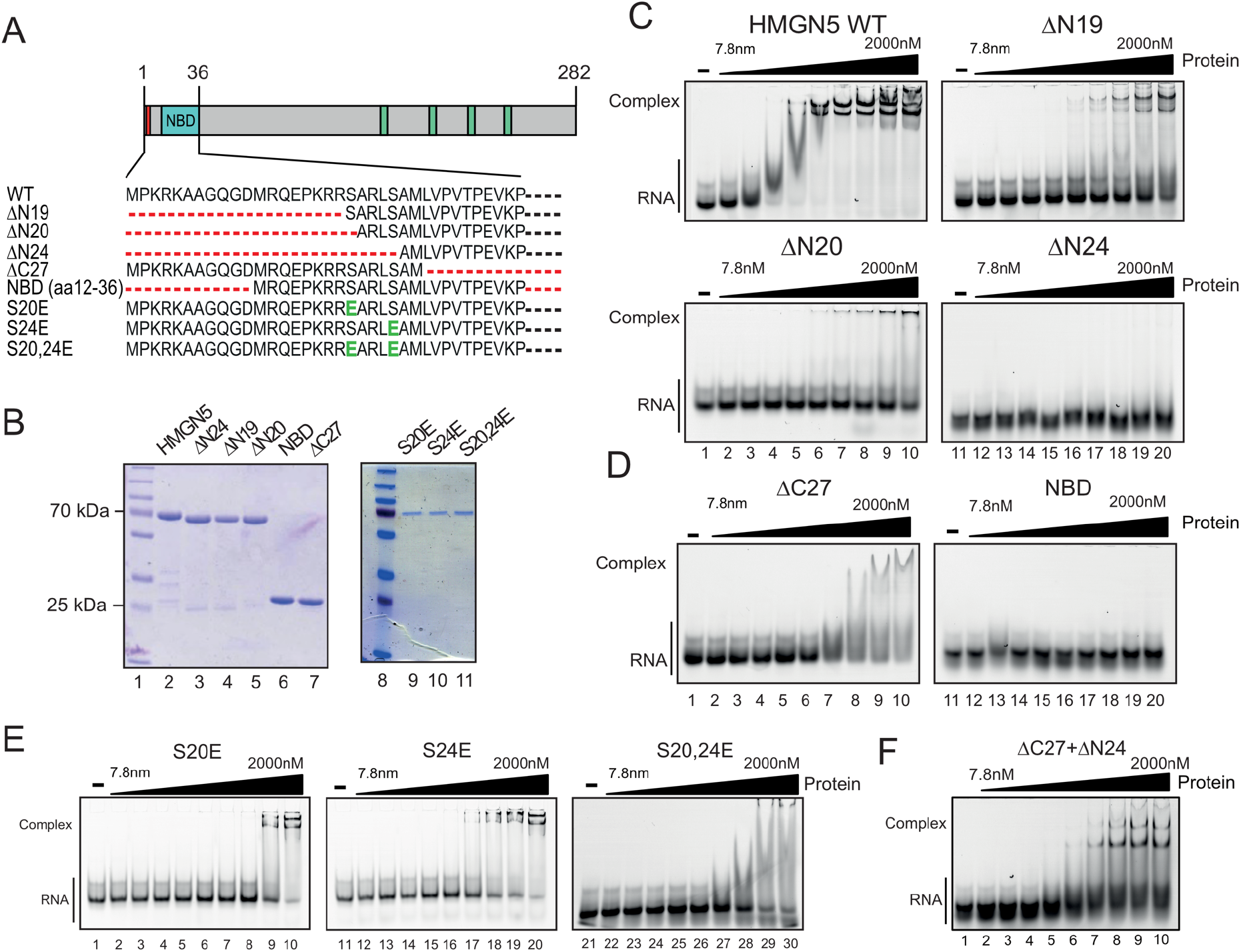
Interaction of deletion and phosphomimetic mutants of HMGN5 with RNA. (A) Schematic representation of HMGN5 deletion and phosphomimetic mutants. The amino acids 1-36 including the nucleosomal binding domain are shown. Red lines represent the deleted amino acids; black lines correspond to not shown amino acids. The phosphomimetic point mutations S20E, S24E and S20,24E are colored in green. (B) Coomassie stained polyacrylamide gel of the recombinant GST-tagged mutants. 1μg of each purified protein was loaded on 12% SDS-PAGE. The 70kDa and 25 kDa bands of the pre-stained protein marker (PageRuler Prestained Protein Ladder Plus) are indicated as molecular weight reference. (C) Interaction of HMGN5 full-length and N-terminal deletion mutants with RNA. Increasing protein concentrations were incubated with the Cy5-labeled snoRNA2T2 and reactions were analyzed by native gel electrophoresis. The RNA was kept at a constant concentration of 50 nM. Proteins were added in increasing concentrations, starting at 7.8nM and increasing 1.5 fold up to 2000nM, as indicated. (D) Interaction of HMGN5 C-terminal deletion mutant Δ C27 and the NBD mutant containing the amino acids 12-36 with RNA. The reaction was analysed as described in (C). (E) Interaction between HMGN5 phosphomimetic mutants and RNA. The phosphomimetic mutants HMGN5 S20E, S24E and S20,24E were analysed as described in (C). (F) Equimolar concentrations of the deletion mutants Δ C27 and Δ N24 were mixed and incubated with RNA. Reactions were analysed as described in (C).

However, the deletion mutant Δ C27 that contains only the first 26 amino acids, including the NBD, did not form a stable RNA-protein complex, exhibiting a strongly reduced RNA binding affinity, when compared to the wild type protein (Figure 2d, Δ C27). Moreover, a peptide representing the full NBD sequence (aa 12-36), did not exhibit any RNA binding, indicating that the NBD sequence is required but not sufficient for RNA binding (Figure 2d). These results were confirmed by analyzing the RNA binding potential of the phosphomimetic point-mutants, which exhibited reduced RNA binding as well (Figure 2e). Mutation of the individual serine residues resulted in a reduction of RNA binding affinity and an additive effect was observed when analyzing the double mutant S20E, S24E (Figure 2e, lanes 27-30). To test for an auxiliary role of the C-terminal domain in RNA binding, we mixed equimolar amounts of the mutants Δ C27 (exhibiting no stable RNA binding) and Δ N24 (a non-binding mutant) and quantified RNA binding by EMSA (Figure 2f). Remarkably, efficient RNA binding activity was observed in EMSA, by simply mixing the two HMGN5 protein fragments (Figure 2f). The result suggests that intramolecular interactions between the NBD and the unstructured C-terminal domain are required for stable and efficient RNA binding.

### HMGN5 binds nascent RNA *in vivo*

To identify potential HMGN5 RNA targets *in vivo* we performed RNA immunoprecipitation after UV crosslinking, using an inducible stable cell line overexpressing GFP-tagged HMGN5 (HMGN5-FlpIn) (Figure 3, Supplementary Figure 2). As control we performed the same experiment with the corresponding inducible cell line expressing the non-modified GFP protein. HMGN5-associated RNAs were isolated using a modified CLIP-seq protocol, which included an additional step of chromatin fragmentation. Isolated RNAs were used for library preparation and Illumina single-end sequencing. We obtained a total of 152,117,704 annotated reads (Supplementary Table 1, Replicates and reproducibility is shown in Supplementary Figure 3). We obtained a total of 6961 significantly enriched peaks, located at 2926 transcripts (Supplementary Table 2). HMGN5 protein was mainly binding to Exons (1887 peaks, representing 29.4%), introns (1751 peaks, 25.1%) and intergenic RNAs (1341 peaks, 19.3%) (Figure 3b and 3c). Performing *de novo* sequence motif analysis, we identified a list of 28 highly enriched RNA binding motifs, of which the top 5 are shown in Figure 3d. Interestingly, amongst the preferential sites we identified motifs enriched in polyA (p-value 1e^-330^) and polyU (p-value 1e^-327^) sequences. These sequences are highly similar to the A-rich (En3RNA_rev) and U-rich (En3RNA) RNAs which were bound by HMGN5 with high affinity *in vitro* (Figure 1b), suggesting that these sequences belong to the specific sequence motifs recognized by HMGN5 correlating with their function *in vivo*.

**Figure 3.**
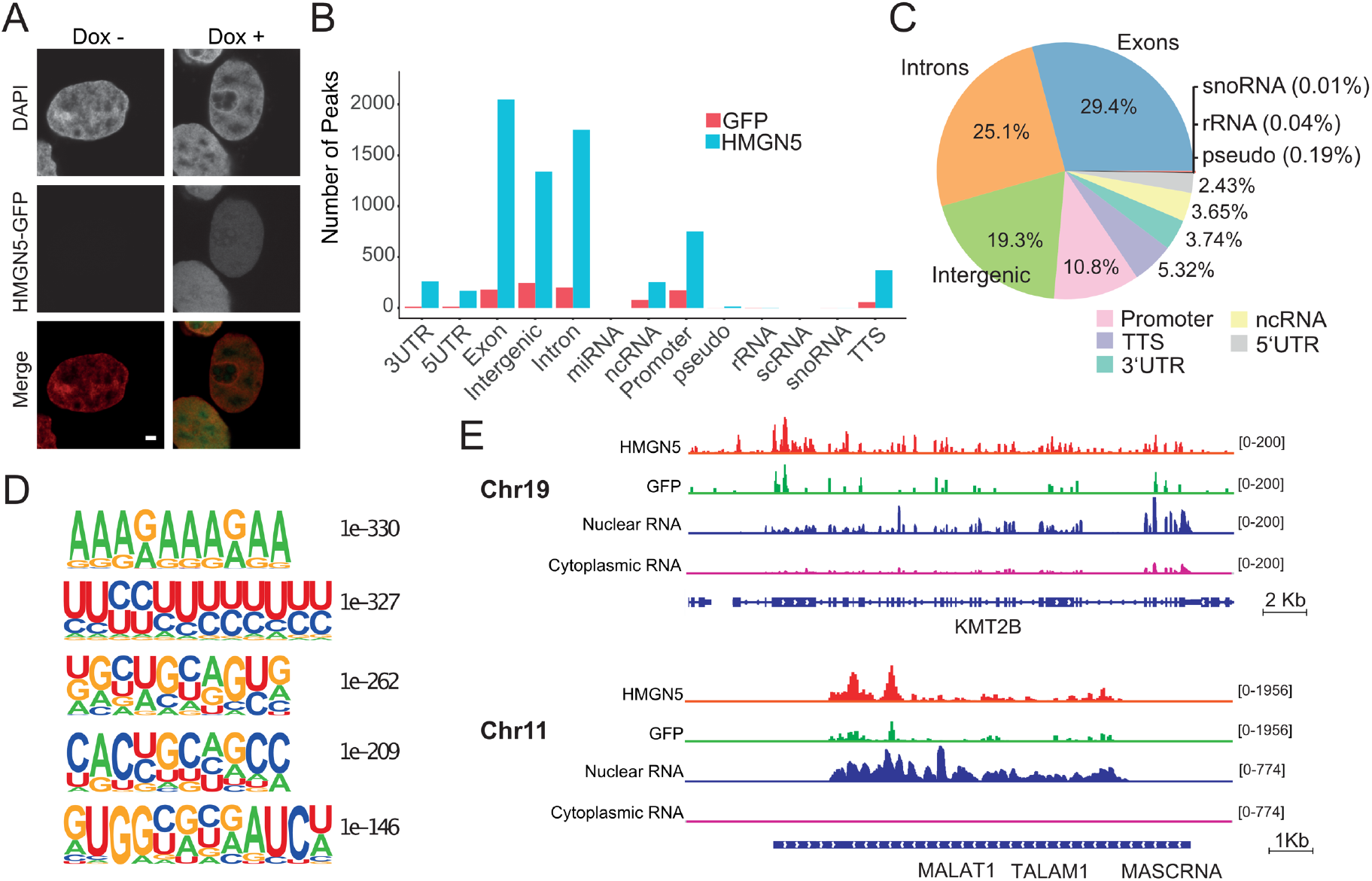
HMGN5 CLIP-seq analysis. (A) Establishment of HMGN5-FlpIn inducible cell line. HMGN5-GFP expression in induced (Dox+) and non-induced (Dox-) HMGN5-FlpIn cells after 24h treatment with or without Doxycycline. Cells were fixed with 4% PFA and analyzed by confocal microscopy on a Leica SP8 device. Scale bar, 2µm. HMGN5 expression was evaluated by GFP fluorescence. DNA was counterstained with DAPI. In merged images, DNA is shown in red and HMGN5 expression in green. (B) Peak distribution HMGN5 CLIP-seq. The number of HMGN5 peaks mapped to each region was compared to those obtained by the control GFP CLIP-seq experiments. The graph represents the number of peaks of HMGN5 in RNAs associated with the selected genomic features. (C) Distribution of HMGN5 peaks represented as percentages. (D) The top 5 most significantly enriched HMGN5 *de novo* motifs. Motifs are shown ranked by p-value. Motif finding was performed with the Homer software using the hg38 reference genome. (E) IGV genome browser tracks depicting representative HMGN5-interacting RNAs. HMGN5 peak track (red) and GFP control (green) are indicated. Distribution of peaks is compared with nuclear (blue) and cytoplasmic RNA (pink) in HEK293T cells. Each track represents the merged alignment from three biological HMGN5 replicates and two GFP biological replicates respectively. KMT2B: Lysine Methyltransferase 2B; MALAT1: Metastasis Associated Lung Adenocarcinoma Transcript 1.

When analyzing the distribution of HMGN5 peaks in the bound RNAs, it is apparent that HMGN5 targets several sites on the same RNA (Figure 3e, Supplementary Figure 4). As examples we show the KMT2B and the MALAT1 RNAs with many HMGN5 binding sites throughout the transcript (Figure 3e). When analyzing the nature of the HMGN5 target RNAs with respect to their subcellular localization, we detected mainly nuclear RNAs, as evidenced by the presence of retained introns (Figure 3e, Supplementary Figure 4). We performed Gene Ontology (GO) analysis of the HMGN5-bound RNAs and found an enrichment of genes associated with RNA metabolic processes (Supplementary Figure 5, Supplementary table 3). Our results suggest that HMGN5 is binding co-transcriptionally to the nascent RNA, indicating a novel role for HMGN5 in regulating the RNA metabolism.

### HMGN5 binds preferentially to active regulatory regions

Next, we analyzed the genomic binding sites of HMGN5 by ChIP-seq analysis, using the inducible HMGN5-FlpIn cell lines. The analysis revealed a total of 8952 specific HMGN5 binding sites (Supplementary Table 4), with a strong enrichment of the protein at introns (35.31% of all peaks), promoters (31.6%) and intergenic regions (24.92%) (Figure 4a). Figure 4b shows a heatmap of the HMGN5 binding profile and the control GFP around the Transcription Start Sites (TSS) of all referenced genes (Ref-seq genes). The analysis shows a specific enrichment of HMGN5 at or close to the TSS. This can be clearly observed in each biological replicate, as shown in detail in Supplementary Figure 6.

**Figure 4.**
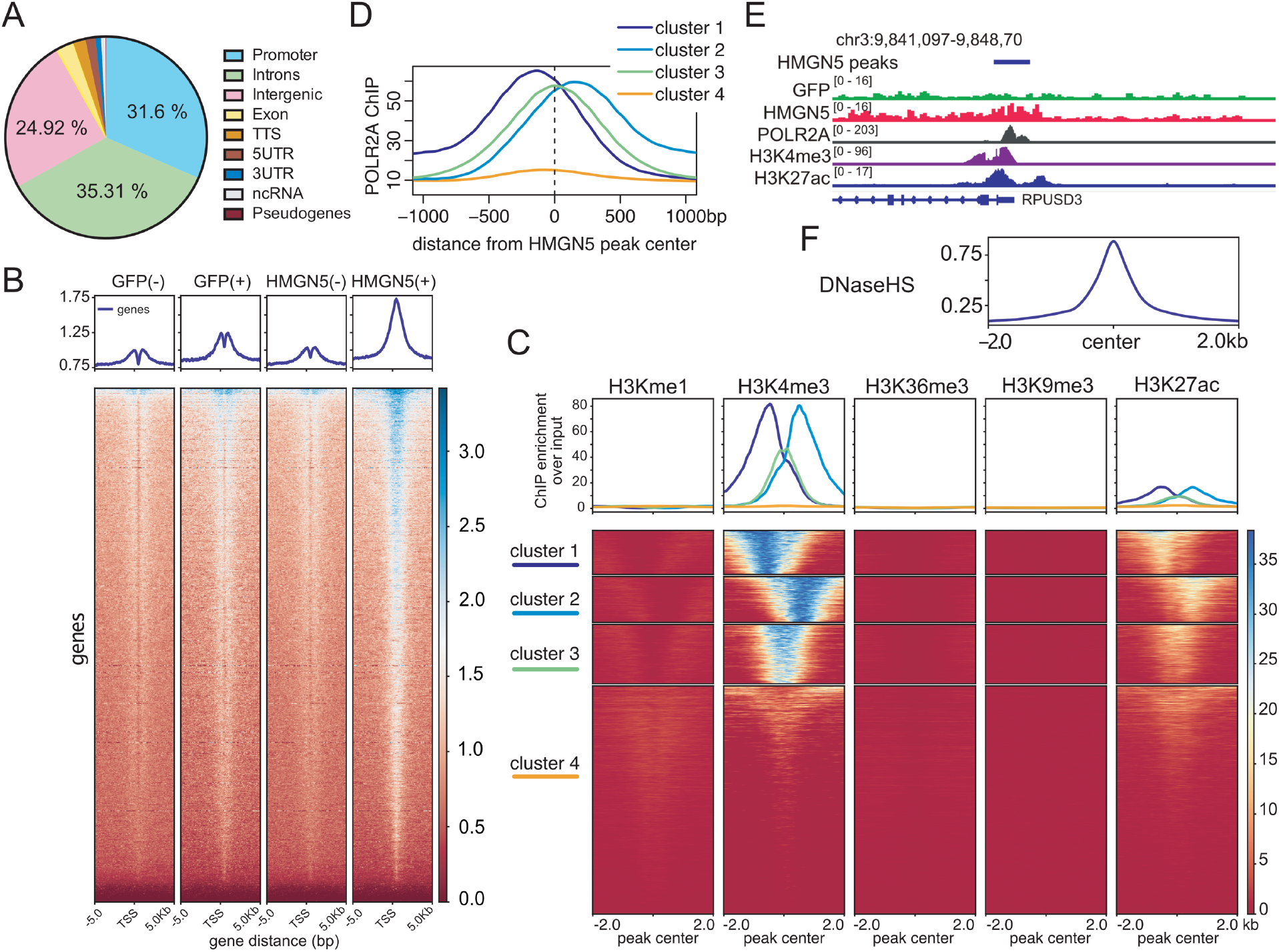
Genome-wide distribution of HMGN5 binding sites. (A) Enrichment of HMGN5 in the selected genomic features. Number of HMGN5 peaks at each genomic attribute is shown as percentage. (B) Heatmap of HMGN5-GFP and GFP control ChIP-seq signals. Genes are ordered vertically by signal strength. GFP(-): uninduced GFP_FlpIn control cell line; GFP(+):GFP-FlpIn control cell line induced with doxycycline; (HMGN5(-): uninduced HMGN5-GFP_FlpIn cell line; HMGN5(+): HMGN5-GFP_FlpIn cell line induced with doxycycline. (C) Clustering of HMGN5 peaks based on histone modifications. Top panel shows the average input normalized ChIP signal of the corresponding histone modification within the cluster. The heatmap illustrates the ChIP signal intensity around each HMGN5 peak. (D) Distribution of RNA Pol II occupancy around HMGN5 binding sites. HMGN5 binding site clusters were derived from histone modification analysis in (C). LOESS (locally weighted scatterplot smoothing) was applied to the raw profile. (E) UCSC genome browser tracks depicting the HMGN5 distribution at an example loci. The genome tracks of the histone marks H3K27Ac, H3K4me3, and RNA Pol II (POLR2A) are also shown. RPUSD3: RNA Pseudouridine Synthase D3. (F) Distribution of DNase-seq signal relative to the center of HMGN5 binding sites.

To correlate HMGN5 binding sites with the activity state of the underlying genomic region, we analyzed the histone posttranslational modifications around the HMGN5 binding sites. For this, we integrated the ENCODE ChIP-seq datasets of histone modifications available for the HEK293 cell line that was used for this study (Figure 4c). We observed that HMGN5 peaks were mainly associated with H3K27ac and H3K4me3 histone modifications, marking active promoter (Figure 4c). This finding confirms the function of the HMGN proteins as chromatin de-condensing factors that are potentially involved in gene activation processes.

However, a detailed analysis revealed that the HMGN5 location relative to the histone marks was not always matching perfectly. Therefore, we grouped the data according to the relative position of HMGN5 peak center to the occurrence of the histone mark, giving rise to 4, clearly distinct clusters. Cluster 1 and 2 indicate an offset between the peak centers, whereas in cluster 3 the HMGN5 and histone mark centers were overlapping (Figure 4c). The differences between cluster 1 and 2 could be easily explained by the different directionality of the genes in the genome. This became evident as well, when analyzing the co-localization of HMGN5 peaks with the presence of RNA Polymerase II. The binding of HMGN5 to promoters of potentially active transcribed genes was confirmed by the identification of a high RNA Polymerase II occupancy at peak clusters exhibiting the active histone marks (Figure 4d). The binding site of HMGN5 is shifted with respect to the direction of transcription (cluster 1 and 2), showing that HMGN5 binds directly upstream of the +1 nucleosome (Fig 4c-d, reverse direction: cluster 1; forward direction: cluster 2).

Gene Ontology analysis of the clustered HMGN5 peaks, indicate an enrichment of genes associated with RNA metabolic pathways for clusters 1-3 (Supplementary Figure 7a) Representative binding of HMGN5 is shown for example loci along with the GFP control, revealing that the binding profiles of the histone marks H3K27ac, H3K4me3 and Pol II correlate with HMGN5 binding (Figure 4e, Supplementary Figure 7b). Furthermore, HMGN5 binding sites do strongly overlap with DNase hypersensitive sites (Figure 4f) ^42^, suggesting that HMGN5 plays a role in chromatin opening at the promoter of active genes. As previously documented ^9^, we observe that HMGN5 induces large-scale chromatin decompaction *in vivo* (Supplementary Figure 7c), when targeted to specific genomic loci. The results show that HMGN5 binds to chromatin at active regulatory regions, allowing chromatin decompaction and potentially regulating gene expression.

### HMGN5 is coupling global chromatin architecture and gene expression

To characterize the role of HMGN5 in transcription regulation, we either overexpressed or knocked down HMGN5 and analyzed global changes in gene expression. Altering HMGN5 expression levels changed the expression of a large number of genes, with 2999 and 3049 genes being affected after HMGN5 overexpression or knock-down, respectively (Figure 5a, Supplementary Table 5). A PlotCount graph of the experiment showed a strict correlation of the expression changes in the biological replicates, showing experimental reproducibility (Supplementary Figure 8). Interestingly, about one third of the gene products (1287) were identified in both, the knock-down and overexpression conditions, indicating a specific function of HMGN5 at these loci. The RNA-seq data was validated by qPCR for four selected candidate genes, being affected by the overexpression and knock-down of HMGN5. Quantitative RT-qPCR was performed on the isolated RNA, showing that the three independent biological replicates matched the RNA-seq results (Figure 5b).

**Figure 5.**
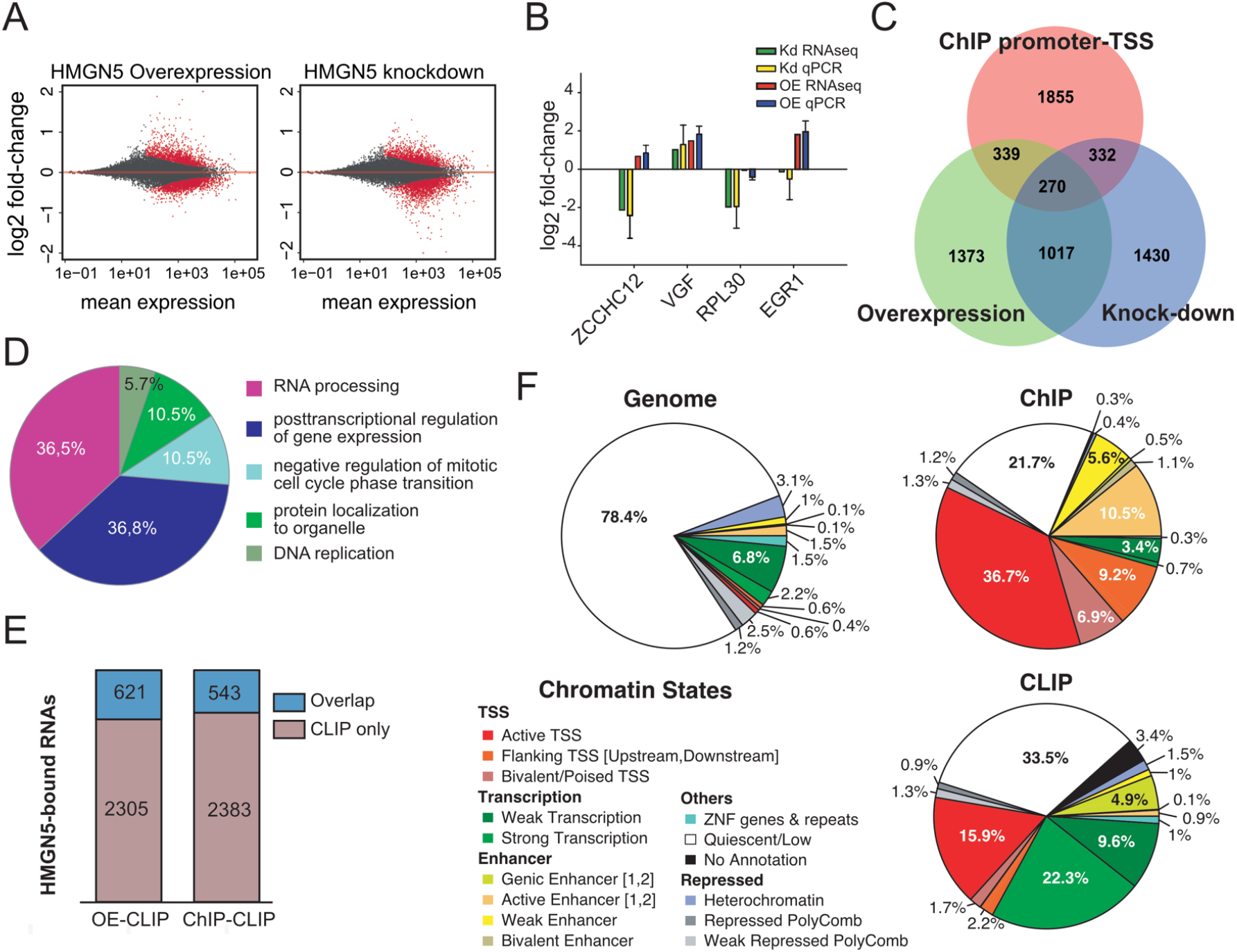
Correlation of HMGN5-dependent transcriptional changes and DNA/RNA binding sites. (A) MA plot of differential expression analysis after HMGN5 overexpression (left) and knock-down (right) compared with the non-treated cells (control). Points representing RNAs with significant differential expression compared with the control samples are marked in red. Gene expression was represented as log2 fold-change ratios. (B) qPCR validation of 4 genes with altered expression. A comparison of log2 fold-change between qPCR and RNA-seq analysis was performed for four selected candidates that showed altered expression between treatments. Standard deviation from 3 independent biological replicates applies only to qPCR results. GAPDH was used for normalization. Two-way ANOVA and Dunett’
ss post-test were used to define statistical differences of mRNA levels between groups. Kd: knock-down; OE: overexpression of HMGN5. ZCCHC12: Zinc Finger CCHC-Type Containing 12; VGF: Nerve Growth Factor Inducible; RPL30: Ribosomal Protein L30; EGR1: Early Growth Response 1. (C) Venn diagram depicting the overlapped set of genes affected after overexpression and knock-down of HMGN5, with the identified HMGN5 binding sites at promoter-TSS. (D) GO analysis of the subset of genes with HMGN5 associated with promoters and showing transcriptional changes after HMGN5 overexpression and knock-down, represented as a pie chart. The analysis was performed with the ClueGo software, and the significance of the enrichment was performed with a hypergeometric test. p-value correction was estimated using the Bonferroni step-down method. The leading term corresponds to the most significant term in the group. Only term with p-value <0.005 are shown. (E) Correlation of HMGN5-associated RNAs found in CLIP-seq, with the RNAs with transcriptional changes after HMGN5 overexpression (OE-CLIP) and with the ChIP peaks (ChIP-CLIP). RNAs found only in CLIP are represented in red, and the overlapped RNAs in blue. (F) Distribution of HMGN5 binding sites along chromatin states of HEK293T cells as defined by the ChromHMM 18-state model. Left pie chart depicts the proportions of chromatin state annotations across the whole genome. Upper (ChIP-seq peaks) and lower (CLIP-seq peaks) pie charts show the proportions of chromatin state annotations assigned to HMGN5 binding sites. Peaks that are not overlapping with any chromatin state annotation were classified as *no Annotation*.

Next, we compared the RNA-seq dataset to the HMGN5 binding sites at gene promoters. We identified a high overlap of 34% (941 peaks) of all HMGN5 binding sites at promoters, to be associated with the genes exhibiting transcriptional changes (Supplementary table 5). This finding indicates a direct functional link between HMGN5 binding and the gene expression changes (Figure 5c). Ontology analysis of the HMGN5 bound promoters, exhibiting differential gene expression in the overexpression and knock-down conditions (270 genes, Figure 5c), showed that these were mainly associated with RNA metabolism (Figure 5d). The most enriched terms were RNA processing and posttranscriptional regulation of gene expression. Intriguingly, the GO analysis of the CLIP-seq data analysis resulted in the same GO terms, suggesting that HMGN5 is at the same time binding to the RNA and the promoters of the genes it is regulating. These results suggest that HMGN5 directly regulates genes involved in the RNA metabolism by targeting their promoter regions and subsequently binds to the emerging RNA.

To quantify the co-binding of promoter and RNA of the same gene, we analyzed the relationship between the genomic HMGN5 binding sites, the differentially regulated genes and the HMGN5-bound RNAs obtained by CLIP-seq (Figure 5e). Interestingly, of the 2926 identified RNA targets, 543 RNAs were found to overlap with genomic HMGN5 binding at promoters, and 622 HMGN5-bound RNAs were affected by overexpression of HMGN5 (Figure 5e, Supplementary table 5). Similarly, when performing GO analysis of these datasets, terms associated with regulation of RNA metabolism appear enriched (Supplementary table 6). Within this overlapping dataset, we did not observe clear correlation between the extent of expression change and HMGN5 RNA/DNA binding strength, however, co-binding is evident for transcriptional changes (Supplementary table 5).

In addition to the strong enrichment of HMGN5 binding sites at gene promoters, we identified a substantial amount of HMGN5 peaks obtained by either CLIP-seq or ChIP-seq in introns or intergenic regions. To test whether HMGN5 might regulate gene expression at distal regulatory regions such as enhancer elements, we analyzed the chromatin states at HGMN5 binding sites using the corresponding ChromHMM 18-state model from ENCODE (Figure 5f). Compared to the genomic background HMGN5 DNA binding sites are highly enriched at enhancer elements (16% of all peaks) and HMGN5 RNA binding sites are enriched at genic enhancer elements (5% of all peaks), respectively.

These results suggest a dual role for HMGN5 in gene regulation. We propose that HMGN5 would first, bind to the target site and open up chromatin by displacing histone H1 ^9^. Second, HMGN5 would be handed over to the nascent RNA, potentially improving overall transcription efficiency or increasing RNA stability.

### HMGN5 forms mutually exclusive complexes with chromatin and RNA

As there is significant overlap between the RNAs bound by HMGN5 and its genomic binding sites, being mostly located in regulatory regions, we asked whether the RNA and nucleosome binding activities in HMGN5 do cooperate. To test this, we performed competitive binding studies using HMGN5 in the presence of RNA and nucleosomes. Specifically positioned nucleosomes were reconstituted on fluorescently labeled DNA, containing the 601 nucleosome positioning sequence ^21^. The Cy3-labeled nucleosomes or the Cy5-labeled snoRNA2T2, were incubated with HMGN5 and analyzed by electromobility shift assays (Figure 6a). HMGN5 binds to nucleosomes, forming a discrete HMGN5-nucleosome complex with low electrophoretic mobility and it also forms a specific complex with RNA (lanes 2 and 3). Both types of HMGN5-complexes can be distinguished, either by their migration properties or the fluorescent labels attached to the substrates. Next, equimolar concentrations of the nucleosomal and RNA substrates were incubated with increasing concentrations of HMGN5 and analyzed (lanes 4 to 10). Evidently, with high HMGN5 concentrations full binding to the RNA and nucleosomal substrate was observed, without the appearance of a detectable HMGN5-RNA-nucleosome complex. Moreover, HMGN5 bound with similar affinity to both substrates, revealing full binding of RNA and nucleosomes at a protein concentration of about 1µM (Figure 6a, lane 9). Next, we tested the stability of the HMGN5-nucleosome complex in presence of RNA. For this, we first titrated the HMGN5 protein to the nucleosome, observing larger HMGN5-nucleosome complexes at high HMGN5 concentrations (Figure 6b). The data suggests that at high HMGN5 concentrations more than one HMGN5 molecule is bound per nucleosome (Figure 6b, lanes 2 to 5). To analyze the effect of RNA on the HMGN5-nucleosome complex, we used an HMGN5 to nucleosome ratio that resulted in the largest HMGN5-nucleosome complex, having more than one HMGN5 protein bound (Figure 6b, lane 6). The addition of excess of RNA did not fully disrupt the HMGN5-nucleosome interactions (Figure 6b, lanes 6-9). We detected a defined, faster migrating HMGN5-nucleosome complex that remained stable, even in the presence of high RNA concentrations (RNA excess ranging between 300ng to 2μg). The experiment suggests a non-dynamic and stable binding of HMGN5 to the nucleosome, albeit the presence of RNA disrupts non-canonical HMGN5-nucleosome or HMGN5 intermolecular interactions. Altogether, these results indicate that HMGN5 exhibits a dual binding mode, either forming a complex with RNA or with the nucleosome. The dual binding mode is supporting the findings of our CLIP and ChIP experiments, proposing a switch from nucleosome to RNA binding during gene activation.

**Figure 6.**
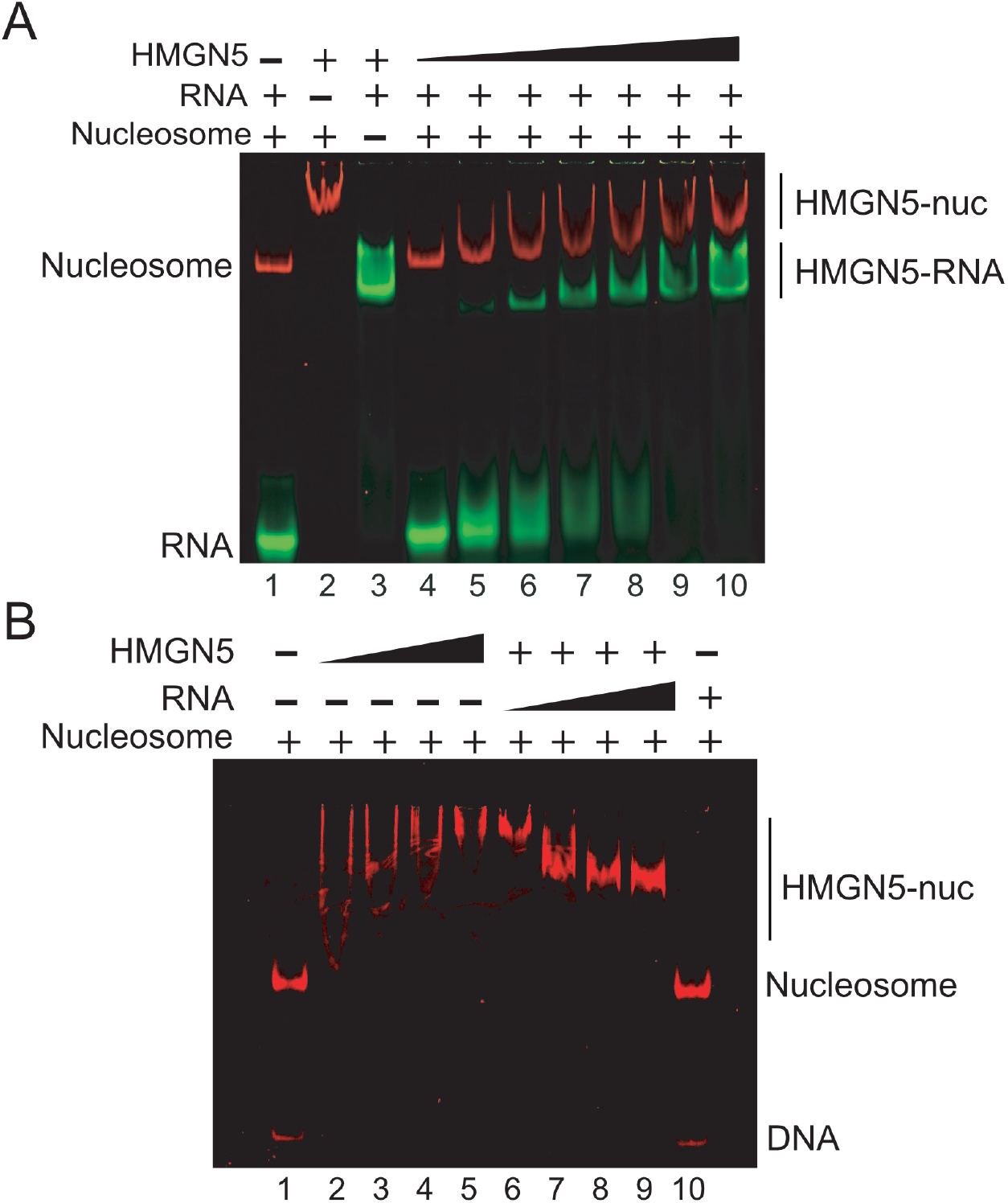
Competitive EMSA showing HMGN5 binding to either RNA or nucleosomes. (A) Competitive EMSA to compare specificity of HMGN5 to nucleosome and to RNA. 50 nM of Cy5 labeled snoRNA2T2 (green) and 50ng of Cy3 labeled mono-nucleosomes (red) were incubated with increasing concentrations of recombinant HMGN5 up to 2 μM (1.5 fold titration). Controls of only RNA+nucleosome (Lane 1), protein+nucleosome (lane 2) and protein+RNA (lane 3) are shown. Nuc: nucleosome. (B) The HMGN5-nucleosome interaction was prepared in presence of an excess of fragmented non-labeled total RNA extracted from HeLa cells. Lane 1: 50ng of Cy3 labeled mononucleosome; lanes 2-5: HMGN5-nucleosome interaction (up to 2 µM protein in lane 5). Lanes 6-9 competition with 300ng, 700ng, 1μg and 2μg of fragmented RNA. Protein and nucleosome were kept constant (2 µM and 50ng respectively). Lane 10, 50ng of mono-nucleosome mixed with 2μg of total RNA. Interactions were loaded on 6% Native PAGE and visualized in a Typhoon 9500 device.

### HMGN5 interacts with CTCF *in vivo*

In order to identify binding partners of HMGN5, we performed immunoprecipitations followed by label-free quantitative proteomics (liquid chromatography-tandem mass spectrometry (LCMS/MS). Since HMGN5 binds tightly to chromatin and to avoid DNA mediated interactions, we treated the cellular lysates with micrococcal nuclease (MNase) and Benzonase to hydrolyze DNA (Figure 7a). Figure 7b shows the affinity purified HMGN5 and the control precipitation of GFP, after the treatment with either MNase and Benzonase, or only with MNase. The immunoprecipitation was efficient in both cases, with lower background when using MNase and Benzonase. The specificity and efficiency of the HMGN5 immunoprecipitation was monitored with the corresponding Western blot (Figure 7c, uncropped Western blot in Supplementary Figure 9a).

**Figure 7.**
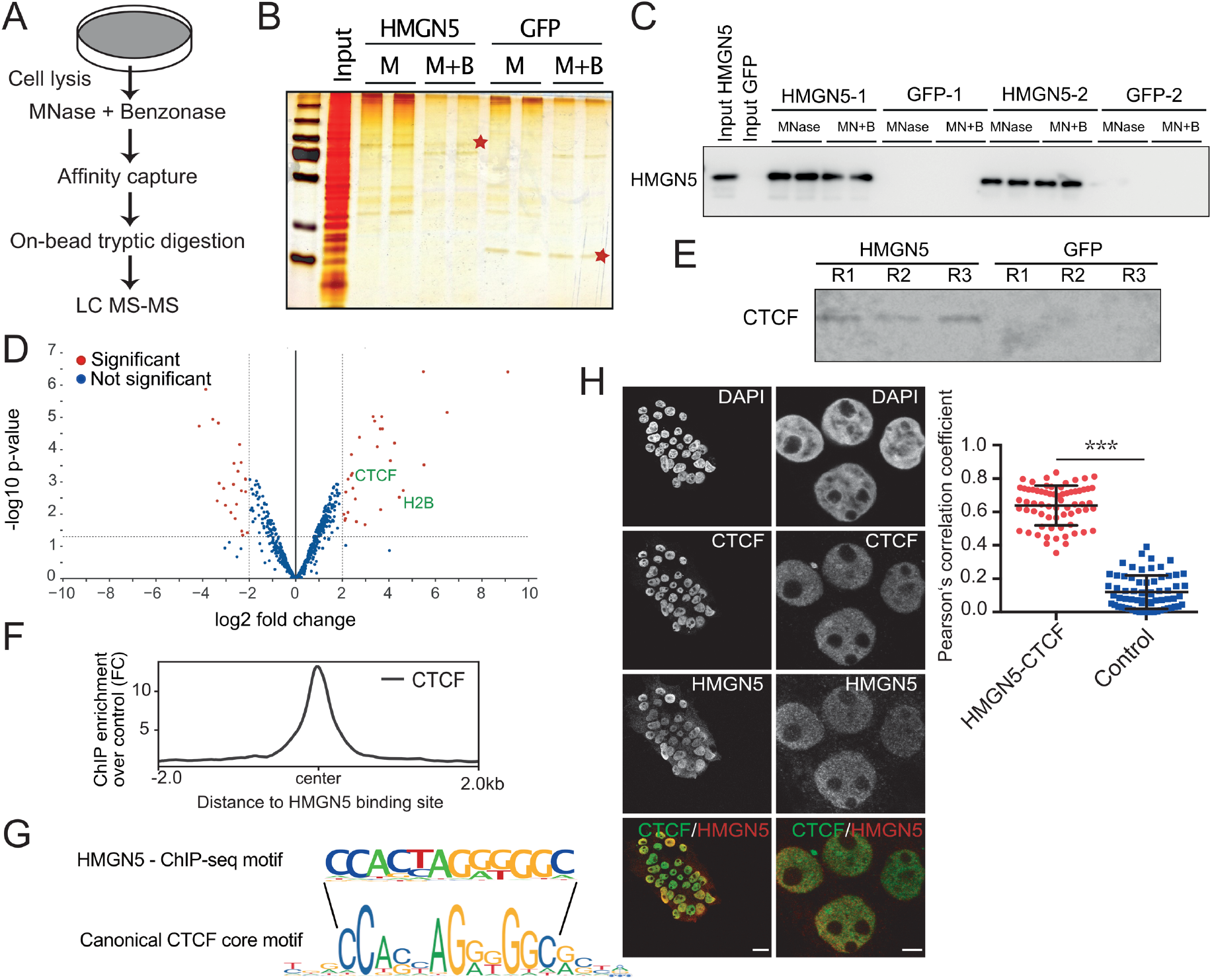
CTCF interacts *in vivo* with HMGN5. (A) Scheme depicting the interactome capture and identification of HMGN5 partners by Co-IP and mass spectrometry. (B) Silver staining after HMGN5 and GFP Co-IP from the FlpIn stable cell lines. M corresponds to MNase digested samples and M+B corresponds to a treatment with MNase plus Benzonase. 10% of the beads used for IP were loaded in each lane. Two biological replicates were loaded per condition. The red stars indicate the overexpressed proteins, HMGN5-GFP (100kDa) and GFP (30 kDa). HMGN5 input was loaded as reference. (C) Validation of the HMGN5 immunoprecipitation by Western blot. Co-IP samples treated with MNase or with MNase plus Benzonase (MN+B) including two biological samples per condition (HMGN5-1, HMGN5-2, GFP-1, GFP-2) and two technical replicates each, were subjected to western blot. Each lane corresponds to 10% of the Co-immunoprecipitated material. (D) Identified HMGN5-interacting partners shown as a volcano plot. Y-axis shows statistical significance -log10 p-value; log2 fold change of HMGN5 over GFP is represented in x-axis. Significantly enriched proteins in the three biological replicates with a log2 fold change ≥2 are shown as red dots. The identified proteins H2B and CTCF are indicated in the plot. (E) Identification of CTCF after HMGN5 immunoprecipitation or GFP control by Western blot. 3 biological replicates (R1, R2, R3) were used per condition. 10% of the Co-immunoprecipitated material was loaded per lane. Antibody sc-28198 (Santa Cruz Biotechnologies) was used for CTCF detection. (F) Correlation of HMGN5 occupancy with the distribution of CTCF binding sites. The average genomic distribution of CTCF was plotted relative to the center of HMGN5 binding sites including a 2000bp window around the peak. (G) Most enriched HMGN5 ChIP-seq *de novo* motif (p-value 1e-141) compared with the canonical CTCF motif obtained from Jaspar database. The lines enclose the canonical CTCF core motif. (H) Co-localization of endogenous CTCF and HMGN5 in HEK293T cells. Left: Confocal images showing the cellular distribution of both proteins. In the merged images CTCF is shown in green and HMGN5 in red. Left panels: 63X magnification (scale 20µm). Right panels: image acquisition using 63X magnification plus 5X digital zoom (scale 5µm). Right: quantitative co-localization by Pearson’s correlation coefficient. Graph represents the mean ± standard error of the mean (SEM), n = 4 biological replicates with a total of 67 cells analyzed by a one-tailed t-test (p < 0.05). Control: co-localization after flipping HMGN5 channel for random co-localization. Double staining for confocal imaging was performed with the CTCF antibody sc-271474 (Santa Cruz) and HMGN5 HPA000511. Nuclei were counterstained with DAPI. Images were taken on a Leica SP8 device.

The quantitative proteomic analysis resulted in the identification of 21 proteins specifically interacting with HMGN5 (Supplementary table 7). Among them, we identified a group of seven proteins involved in the regulation of ribosomal RNA metabolism (RPL10, LAS1, PELP1, TEX10, WDR18, SENP3, NOL9) (Supplementary table 7), these proteins, with the exception of RPL10, form a stable multi-protein complex involved in the pre-rRNA processing and synthesis of the 60S ribosomal subunit ^43,44^.

Furthermore, we identified CTCF as an interacting partner of HMGN5 *in vivo* (Figure 7d, Supplementary table 7) and validated this interaction by Western blot analysis (Figure 7e, uncropped Western blot in Supplementary Figure 9b). CTCF constitutes the major organizer of chromatin architecture in the nucleus, having a pivotal role in establishing the 3D genome organization and functioning as an insulator ^45^. The association of CTCF with HMGN5 suggests a joint role of both proteins in the organization of higher order structures of chromatin. Indeed, when analyzing the genomic occupancy of HMGN5 compared to CTCF in the same cell line (HEK293; ENCODE accession ENCFF128UTY), we observed that CTCF binding sites do overlap with the HMGN5 peak center, identified in our ChIP-seq experiments (Figure 7f). A *de novo* motif analysis, using the HMGN5 ChIP-seq dataset revealed that the most enriched HMGN5 motif corresponded to the canonical CTCF binding site ^46,47^ (Figure 7g; see Supplementary Figure 10a for all enriched motifs). The CTCF motif was identified in 17 percent of all genomic HMGN5 binding sites, indicating a high co-occupancy of both proteins. Such a high co-occupancy is also evident when analyzing the co-localization of endogenous HMGN5 and CTCF by confocal microscopy (Figure 7h). The co-occupancy anaylsis, performed by pearson’s correlation coefficient, indicates that both proteins present a “strong” correlation in cells, according to the co-localization degree for Pearson’s ^23^ with a mean coefficient of 0.63, which was significantly higher compared with the control of random co-localization events (co-localization with flipped HMGN5 channel), which showed correlation values close to 0.

To test whether HMGN5 on its own recognizes the DNA binding site of CTCF, or whether HMGN5 is recruited to these motifs sites via its interaction with CTCF, we performed DNA electromobility shift assays (EMSA). Increasing concentrations of HMGN5 were incubated with a fluorescently labeled CTCF DNA binding motif or its mutated form in a competitive EMSA assay (Supplementary Figure 10b). Mixing the Cy5 labeled CTCF DNA with the Cy3 labeled mutated CTCF binding site showed that HMGN5 binds to DNA at high concentration but is not specifically binding to the CTCF motif. The results suggest that HMGN5 is recruited via CTCF to its genomic binding sites.

In summary, these results hint to an important role of HMGN5 in regulating the accessibility of specific CTCF-domains, maintaining an open chromatin conformation by displacing factors like H1, opening the regulatory elements and subsequently binding the newly synthesized transcripts to regulate transcription.

## Discussion

HMGN5 is a member of the HMGN family that regulates gene expression by inducing large-scale chromatin de-condensation, by directly interacting with the nucleosome ^9,10,12^.

We previously showed that nuclear RNA associates with chromatin to keep its structure accessible and transcriptionally active ^18^. Our laboratory demonstrated that in *Drosophila*, an RNP complex formed by chromatin-associated snoRNA together with the chromatin-associated protein Df31, was required for the opening and maintenance of higher order structure of chromatin ^18^. In humans, the protein HMGN5 shares common functional and structural features with Df31, such as the ability of chromatin de-compaction and its intrinsically disordered structure and negatively charged amino acids. However, the molecular mechanism by which HMGN5 is targeted to its site of activity and how it regulates the changes of chromatin structure is not fully understood.

Here we show for the first time that HMGN5 possesses a novel and specific RNA binding activity, the protein exhibits a strong preference for binding ssRNA over ssDNA of the same sequence. Remarkably, our data shows that the RNA binding domain overlaps with the NBD domain, which mediates the binding to nucleosomes, and this binding is stabilized by the negatively charged C-terminal region of HMGN5, revealing the necessity of two protein domains and their intramolecular interaction for stable RNA binding. This feature has been previously described for other proteins containing intrinsically disordered domains ^48^. Moreover, the latest evidence suggests that intrinsically disordered regions, as in the case of HMGN5, contribute to RNA binding ^49,50^. Intrinsically disordered proteins may mediate phase separation and the establishment of specialized nuclear compartments, as shown of SAF-A through interaction with nuclear RNA ^51,52^.

RNA is an active component of complexes and proteins regulating chromatin structure, such as HMGA1 ^53^, YY1 ^54^, or CTCF ^55,56^. Here, we provide the first evidence of RNA binding to a member of the HMGN family, the only known proteins that change chromatin architecture by directly interacting with the nucleosome ^57^. Binding to RNA is extended to other members of the HMGN family, as HMGN2 was also able to specifically interact with RNA *in vitro*, suggesting a functional specificity of RNA binding for HMGN.

Our CLIP-seq study, showed that HMGN5 is binding co-transcriptionally to the nascent RNAs, as similar levels of HMGN5 binding to exons and introns was observed, (29% and 25% of all peaks, respectively), suggesting that HMGN5 exerts a co-transcriptional regulation. Moreover, the identification of specific consensus motifs in the HMGN5-bound RNA highlights that the binding is sequence-dependent, which is supported by our *in vitro* data. Interestingly, HMGN5-bound RNAs are mainly clustered in pathways associated with RNA metabolic processes, highlighting a role for HMGN5 in regulating RNA metabolism.

Additional ChIP-seq experiments revealed that HMGN5 did not only bind to nascent RNA but is preferentially associated to regulatory regions of the genome, including active promoters and enhancers (Figure 4 and 5). Our data suggests that HMGN5 binds upstream of the +1 nucleosome to active promoter, which are associated with active histone marks, high RNA polymerase II occupancy and DNase hypersensitive sites. These facts do clearly reveal an association of HMGN5 with active transcription.

A similar genomic profile has been shown for HMGN1 and HMGN2, demonstrating that those HMGN variants are required to maintain the DHS landscape and open chromatin architecture in several mammalian cell lines ^58-62^. A functional redundancy of HMGN5 with other HMGN members, like HMGN1 and HMGN2, can be assumed ^60^.

A large fraction of the HMGN5 ChIP peaks at promoter regions (34%) were associated with changes of gene expression of the same gene, when overexpressing, or knocking down HMGN5. The results implicate a direct functional, mechanistically linking HMGN5 binding to the regulation of gene expression. Furthermore, GO analysis of these genes, resulted in the same GO terms as found for the HMGN5 bound RNAs, as determined by the CLIP-seq experiment. Both, regulated genes and bound RNA are described by the GO term RNA metabolism, implying the co-occupation of the gene promoter and subsequently as well the RNA generated by this locus. An overlap that we could directly show. Altogether, these findings suggest that HMGN5 regulates RNA metabolism by a dual binding mechanism. First, opening chromatin structure at regulatory genomic regions, and second, a co-transcriptional regulation by directly contacting nascent RNA. HMGN5 could either stabilize the pre-mRNA or enhance transcription rates by binding.

The function of HMGN5 in RNA metabolism is further strengthened by the fact that we found HMGN5 being associated with seven proteins (RPL10, LAS1, PELP1, TEX10, WDR18, SENP3, NOL9) that regulate ribosomal RNA metabolism (Supplementary table 7), 6 of them part of a protein complex involved in the pre-rRNA processing and synthesis of the 60S ribosomal subunit ^43,44^.

To obtain more mechanistic details of this potential two-steps mechanism, we performed *in vitro* RNA-nucleosome competition assays (Figure 6). To our surprise, HMGN5 was able to exclusively interact either with the nucleosome or with RNA. Therefore, we suggest that HMGN5 cannot bind to its genomic locus and the pre-mRNA at the same time. Therefore, we suggest that HMGN5 is first bound to the promoter region upstream of the +1 nucleosome, de-compacts chromatin, cooperates in gene activation and then is handed over to the nascent RNA to fulfill additional functions that increase the cellular transcript level. We envision a switch from a chromatin bound to an RNA bound form of HMGN5, or alternatively the additional recruitment of HMGN5 to be loaded on the RNA during gene activation. Such a dual function has been shown for EZH2, the catalytic subunit of the Polycomb repressive complex 2 (PRC2), in a mechanism by which PRC2 senses the activation states of promoters through contacting nascent RNAs ^63^. It was later demonstrated that PRC2 forms mutually exclusive complexes with either chromatin or with RNA ^64^, similar to our HMGN5 results. A central question that remains unanswered is whether RNA is required for the recruitment of HMGN5 to its genomic target sites, and if RNA is required for chromatin de-condensation. We hypothesize that HMGN5 may form distinct interaction complexes that could mediate the recruitment to its target sites. This effect could be mediated by un-known regulatory factors, such as proteins or regulatory RNAs.

Our results indicate that one third of the genes affected by HMGN5 de-regulation were overlapping between HMGN5 up and down regulation, showing the same type of change. In contrast to the expectation, most of these genes are repressed in the absence of HMGN5, and also when HMGN5 is overexpressed. We suggest that HMGN5 is part of a multifactor complex (or complexes) and the depletion or overexpression of HMGN5 may produce an imbalance in stoichiometry and disrupting the functional complex. As a result, the same phenotype would be observed in both conditions. Analogous phenotypes with depletion and overexpression of proteins have been observed in stoichiometrically balanced complexes ^65-68^.

Besides the known interaction with the linker histone H1, HMGN5 has been shown to interact with the Lamina-associated Polypeptide 2 α (LAP2α) ^69^, and with Hsp27 ^70^. Here, we identified 21 novel HMGN5-protein partners by quantitative mass spectrometry. Among them, we identified CTCF, a pivotal chromatin architectural protein required for the 3D genome organization ^71^. We revealed that HMGN5 co-localizes with CTCF in cells and physically interacts with the protein *in vivo*. The ChIP-seq data revealed that 17% of total HMGN5 peaks corresponded to the canonical CTCF binding site. The strict correlation that we observe with CTCF binding sites and the enrichment of HMGN5 in regulatory regions of genes, highly suggests that a subpopulation of both proteins may cooperate in the same biological process. Similarly, it was previously shown that HMGN1 binds genome-wide to the edge of nucleosomes surrounding CTCF in CD4+ T cells ^59^, opening the possibility of a common regulatory mechanism for the HMGN family. The inability of HMGN5 to specifically recognize the DNA binding motif of CTCF, suggests that its location at CTCF binding sites in the genome are due to a direct interaction of both proteins. Our results hint to a mechanism in which HMGN5 may open chromatin at specific CTCF-domains, by H1 displacement and being afterward transferred to the nascent transcript to regulate transcription.

## Supplemental information

Supplemental Information includes 12 Figures and 10 tables.

## Acknowledgements

We thank Dr. Attila Németh for critical discussions; Dr. Julia Wimmer for her support with High throughput analysis; Elisabeth Silberhorn, Dr. Helen Hoffmeister and Dr. Rodrigo Maldonado Águila for experimental support and suggestions; Ignasi Forne from Axel Imhof’s group for mass spectrometry analysis; Armando Amaro for assistance with production of the figures; Dr. Andrea Bleckmann for support with image acquisition. This work was supported by the project SFB960 from the Deutsche Forschungsgemeinschaft.

ANID/Fondecyt 11190998 and ANID-Basal funding for Scientific and Technological Center of Excellence, IMPACT, FB210024.

## Author contributions

Conceptualization & experiments design, G.L. and I.A.; I.A. performed most of the experiments.; G.N. performed CLIP-seq ChIP-seq and RNA-seq bioinformatics; I.C. performed the RT-qPCR; U.S. supported RNA-seq analysis and performed the correlation analyses with ENCODE data; S.B. performed EMSA experiments. M.B. performed quantitative co-localization analysis. Mass spectrometry was performed in collaboration with A.I.; G.L and I.A. wrote and edited the text.; M.M. critically read the manuscript and supported text editing. Funding Acquisition, G.L.

