## Supplementary for "HMGN5, an RNA or Nucleosome binding protein - potentially switching between the substrates to regulate gene expression"

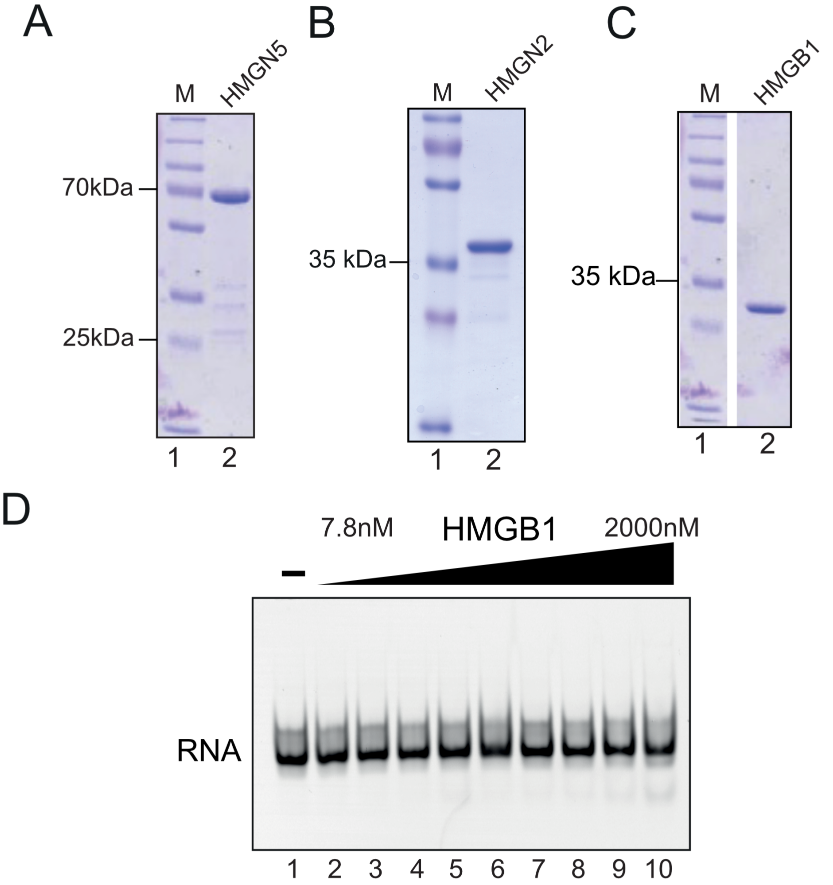


**Supplementary Figure 1**. Recombinant proteins and analysis of HMGB1-RNA interaction. (A - C) Coomassie blue stained gels of the purified, recombinant, GST-tagged HMGN5 protein, HMGN2 and his-tagged HMGB1, as indicated. The purified protein (1µg) was analyzed by SDS-PAGE (12% polyacrylamide). The pre-stained protein marker (PageRuler Prestained Protein Ladder Plus) was used as molecular weight reference (M). (D) Interaction of HMGB1 with RNA analyzed by EMSA. The Cy5-labeled RNA snoRNA2T2 was kept at a constant concentration of 50nM (lanes 1 to 10). HMGB1 was used with an increase of 1.5 fold concentration steps, starting a 7.8nM to 2000nM (lanes 2 to 10). Reactions were analyzed by native gel electrophoresis on 6% polyacrylamide gels in 0.4x TBE buffer and documented with the fluorescence imager Typhoon FLA 9500 (GE Healthcare).

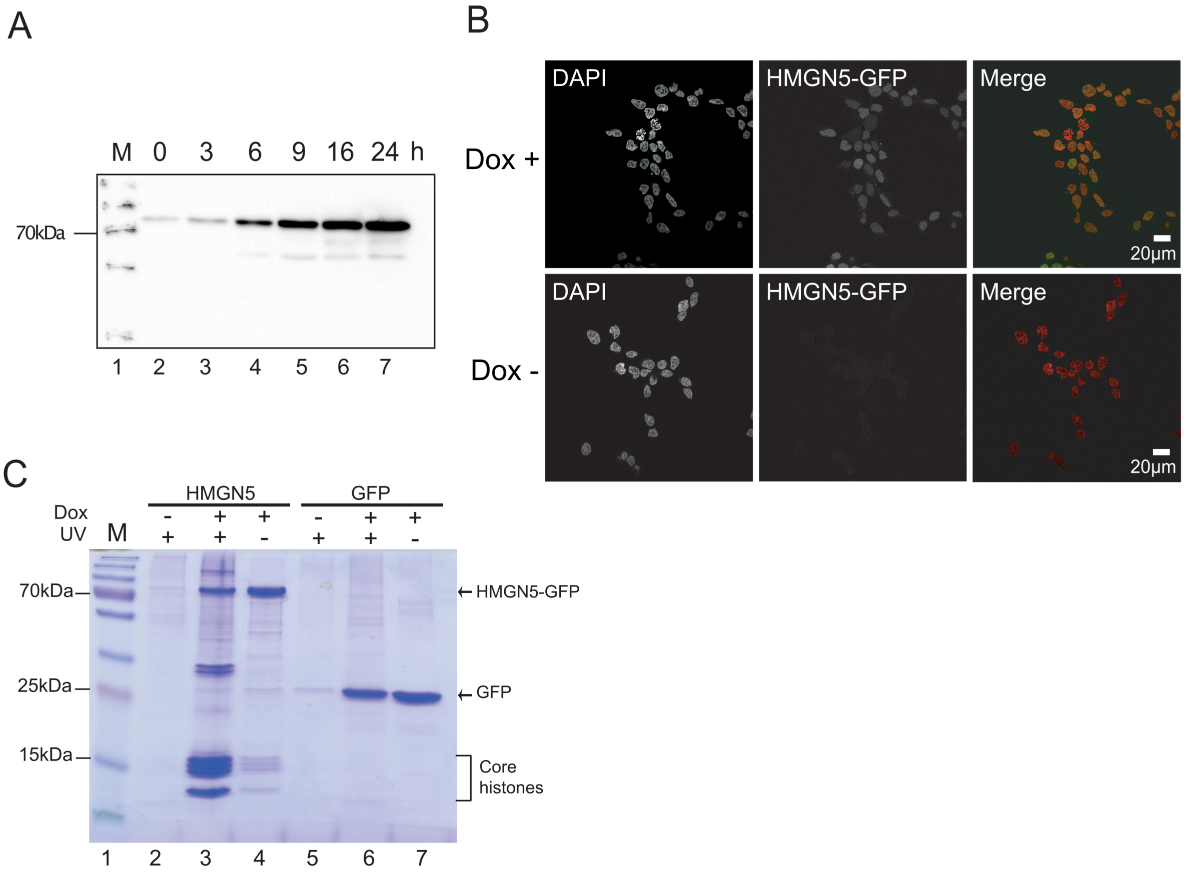


**Supplementary Figure 2.** Establishment of inducible HMGN5-FlpIn cell lines and testing the efficiency of UV-crosslinking. (A) Time-resolved expression of HMGN5-GFP after induction by Doxycycline (1 µg/ml; time points are indicated), using a clonal HMGN5-FlpIn cell line. 20µg of total protein were subjected to immunoblotting, using the anti-HMGN5 antibody HPA000511 (Sigma). The 70kDa band of the protein marker (M) is indicated. (B) The stably transformed HMGN5-GFP cell line was seeded in 6-well plates. After 24h of induction with (Dox+) or without Doxycycline (Dox-), cells were fixed with 4% PFA and analyzed by confocal microscopy (Leica SP8). DNA was stained with DAPI and the HMGN5 expression was visualized by GFP fluorescence. The merged images show the DNA in red and HMGN5 in green. The scale bar measures 20µm. (C) UV-crosslinking conditions used for the experiments. Cells grown in 15cm plates (HMGN5-FlpIn (lanes 2 to 4) and GFP-FlpIn cell lines (lanes 5 to 7)), were treated with or without Doxycycline (Dox), and subjected to UV-crosslinking by applying 150mJ/cm$^2$ UV at 254nm (UV+), or without UV treatment (UV-), as indicated. Protein purification was performed with GFP_Trap agarose beads to reveal interacting proteins by SDS-PAGE. 10% of the immunoprecipitated samples were loaded on a 12% SDS-PAGE and visualized after Coomassie staining. The PageRuler Prestained Protein Ladder Plus was used as molecular weight marker (M).

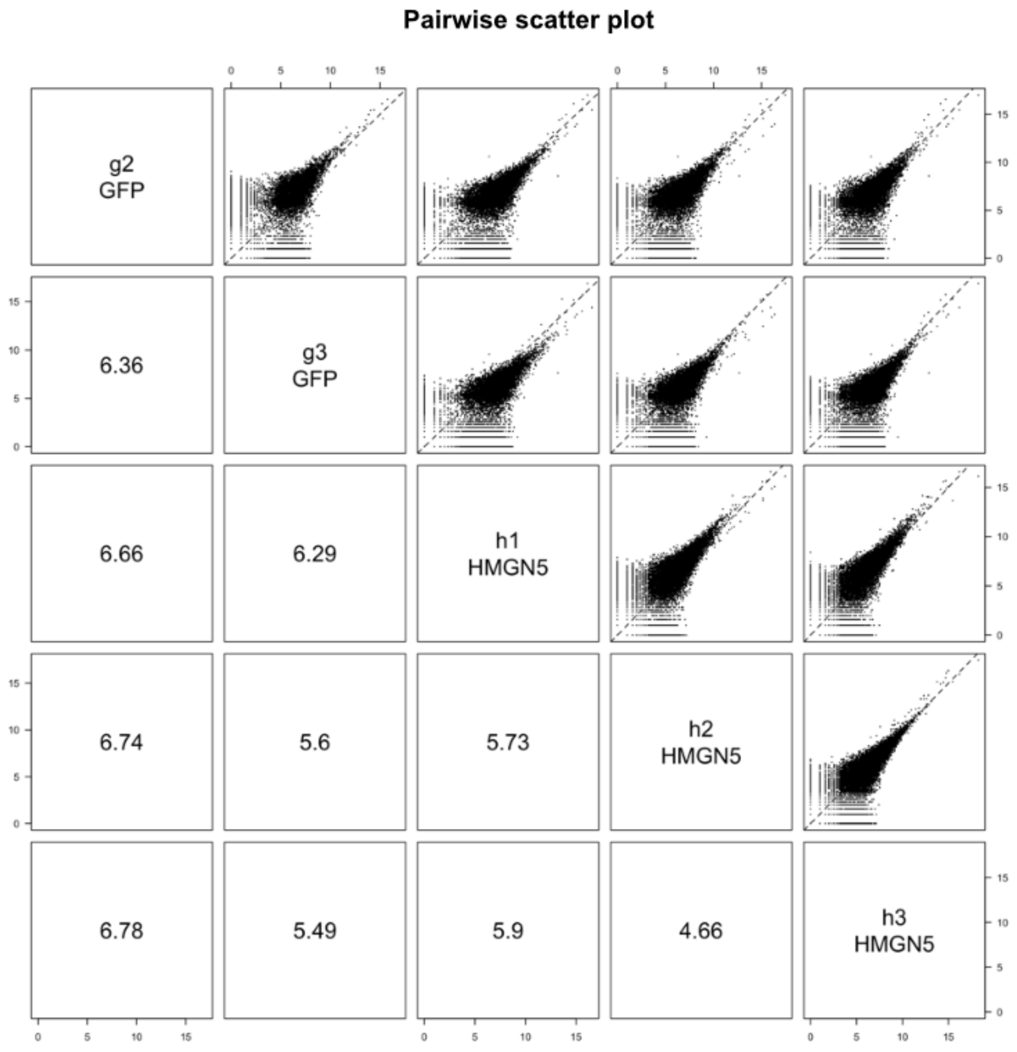


**Supplementary Figure 3.** Pairwise comparison of experimental CLIP-seq samples. Pairwise Scatter plot from HMGN5 and GFP biological replicates. The data is represented as $\log_2$(counts+1).

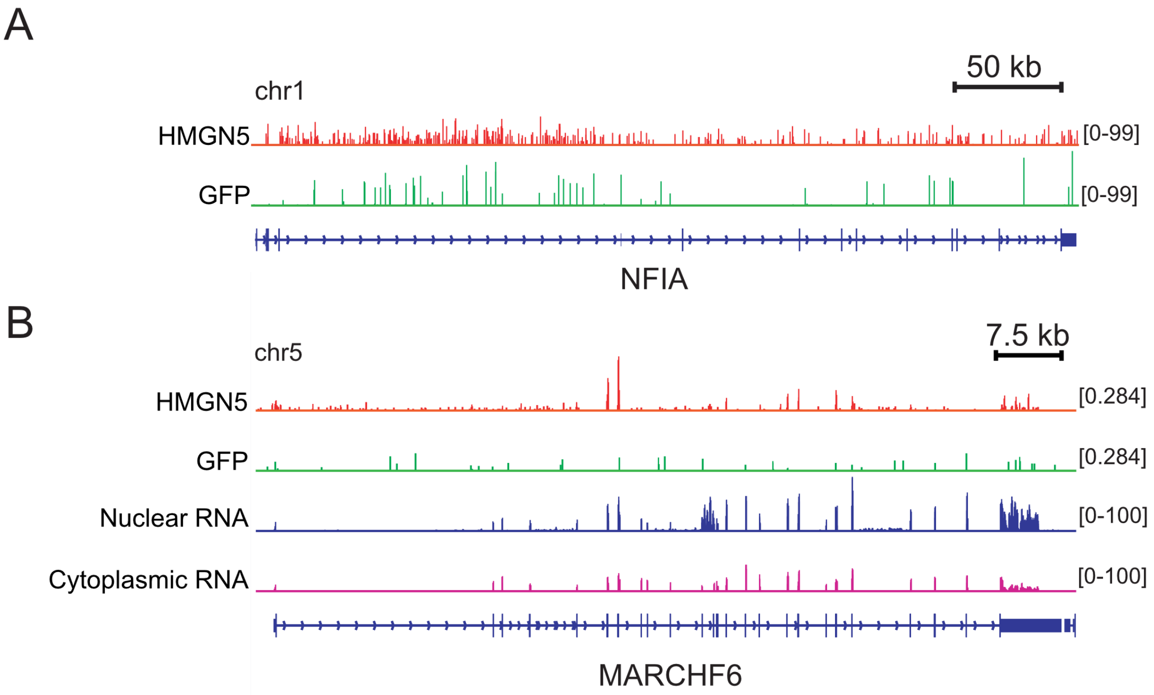


**Supplementary Figure 4.** Extension of the data presented in Figure 3e. HMGN5 peak tracks (red) and GFP controls (green) are indicated. (A) View on the genomic region NFIA: nuclear factor 1 A-type. (B) View on the genomic region MARCHF6: Membrane Associated Ring-CH-Type Finger 6. For MARCHF6, the analysis of nuclear and cytoplasmic RNA are also shown.

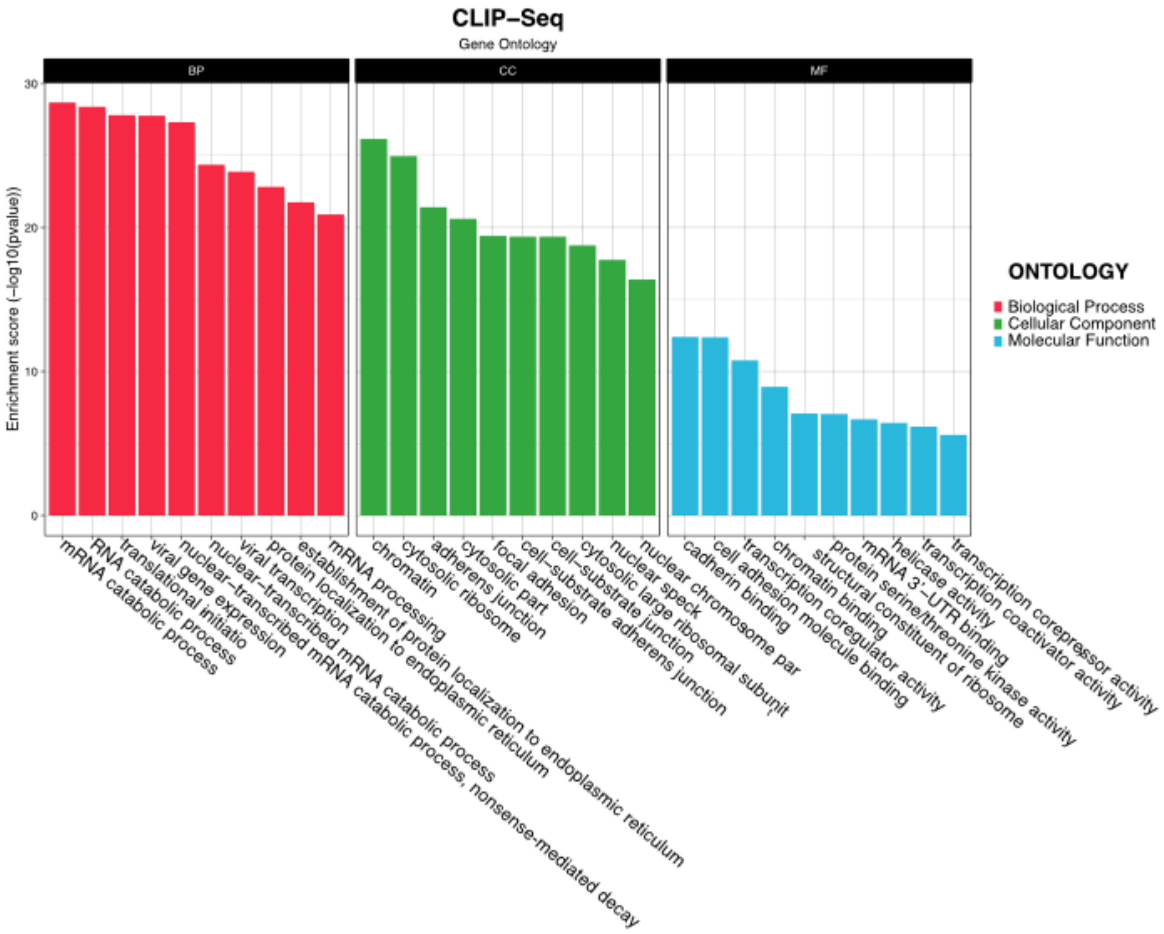


**Supplementary Figure 5.** GO analysis of HMGN5-bound RNAs.

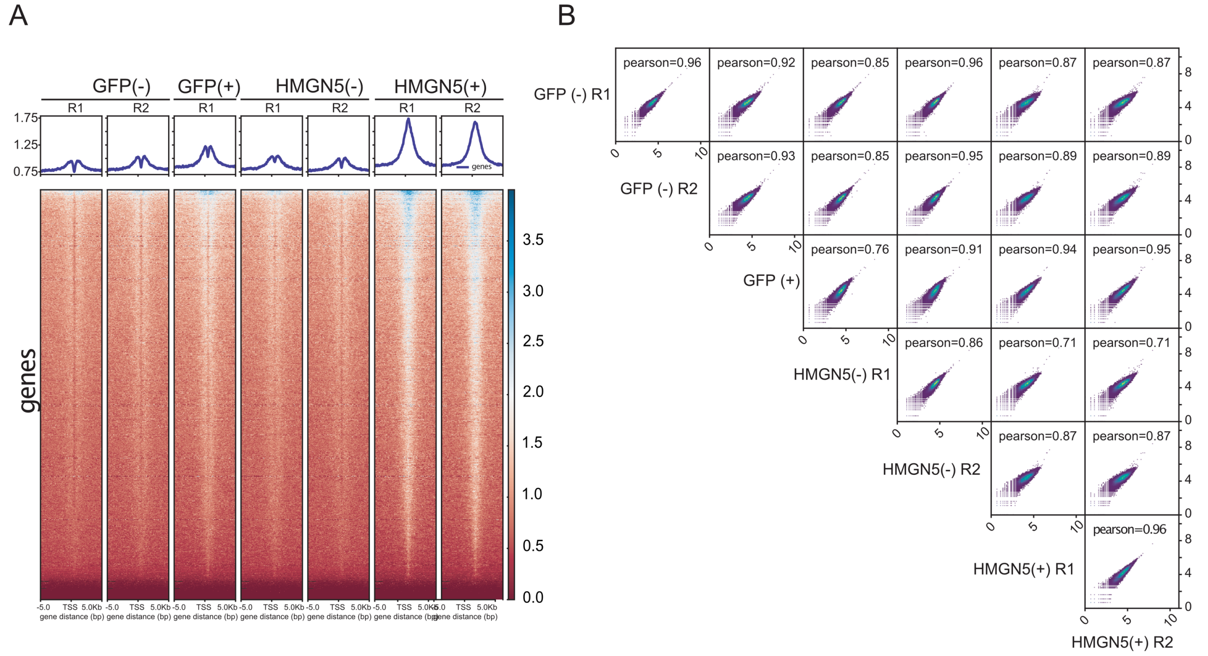


**Supplementary Figure 6.** Reproducibility of the ChIP-Seq experiment. (A) Heatmap of HMGN5-GFP and GFP control ChIP-seq signals of the independent replicates, plotting binding signals around the TSS of Ref-Seq genes. Peak calling was performed for each replicate independently. Genes are ordered vertically by signal strength. GFP(-): uninduced GFP_FlpIn control cell line; GFP(+): GFP-FlpIn control cell line induced with doxycycline; HMGN5(-): uninduced HMGN5-GFP_FlpIn cell line; HMGN5(+):HMGN5-GFP_FlpIn cell line induced with doxycycline. R1 and R2: independent biological replicates. (B) Pairwise Scatter plot of the HMGN5 and GFP biological replicates. The data is represented as $\log_2$(counts+1).

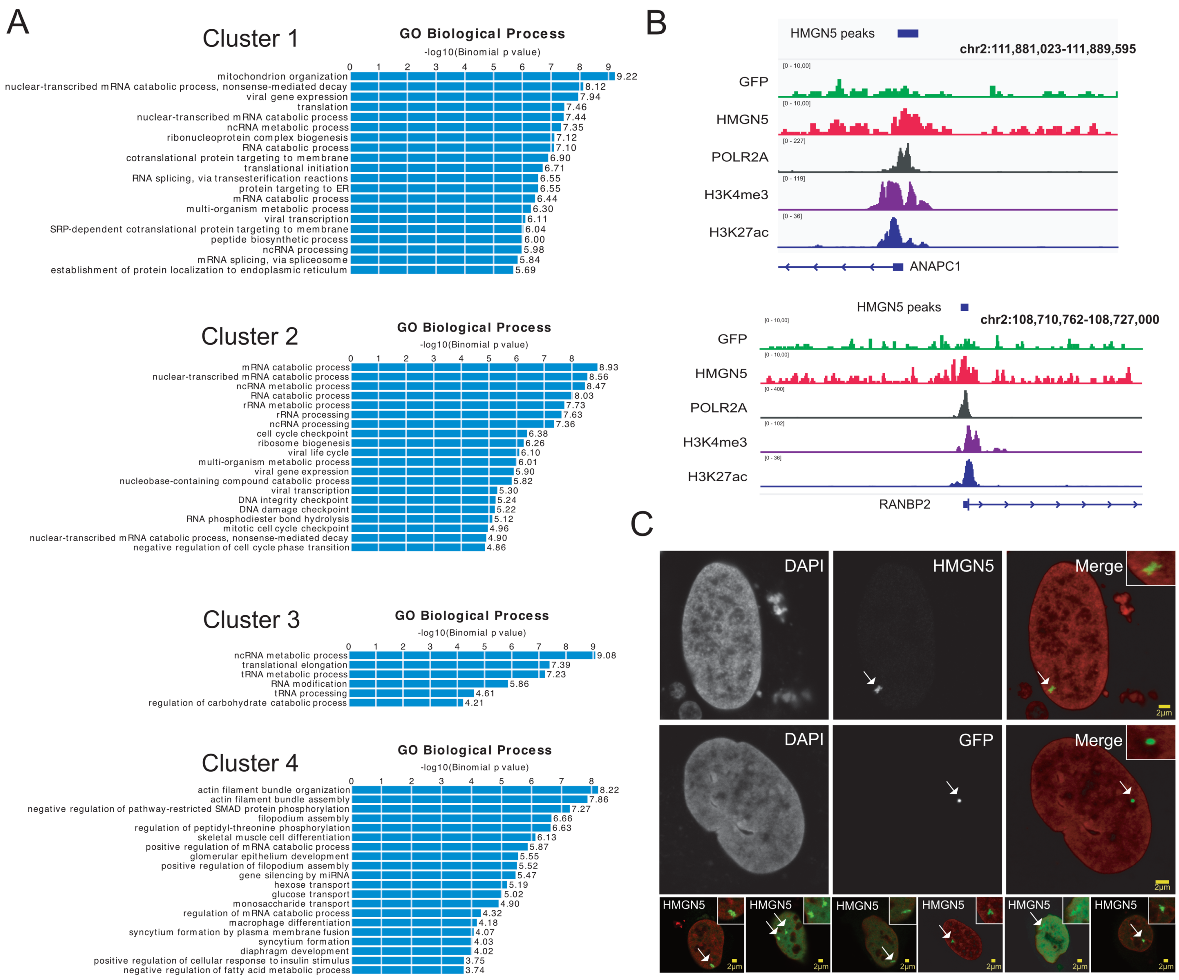


**Supplementary Figure 7.** (A) GO analysis of clustered HMGN5 ChIP peaks from Figure 4c and 4d. GO was performed with GREAT version 4.0.4 (http://great.stanford.edu/public/html/index.php). (B) Genome browser snapshot of HMGN5 binding at 2 different loci. ANAPC1: Anaphase-Promoting Complex Subunit 1; RANBP2: RAN binding Protein 2. (C) Chromatin de-compaction analysis by LacI/LacO tethering. Cells were transfected with the plasmids pSV2_GFP_LacI_HMGN5 or pSV2_GFP-LacI for transient expression of HMGN5-GFP-LacI or the control GFP-LacI, respectively. Chromosomal tethering of HMGN5 was followed by GFP signal. Insets show magnified areas indicated by the arrows. In the merged images, DNA is shown in red and GFP signal in green. Scale bars represent 2µm. Immunofluorescence acquisition was performed on a Leica SP8 microscope.

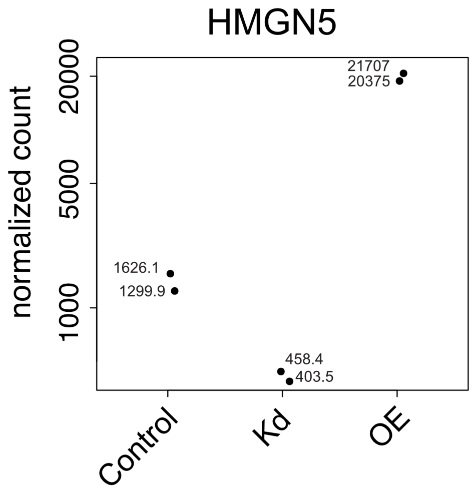


**Supplementary Figure 8**. PlotCount graph showing the normalized count of the HMGN5 transcripts from the RNA-seq in the non-treated (Control), knock-down (Kd) and overexpression (OE) conditions of HMGN5. Each dot in the groups represents a biological replicate.

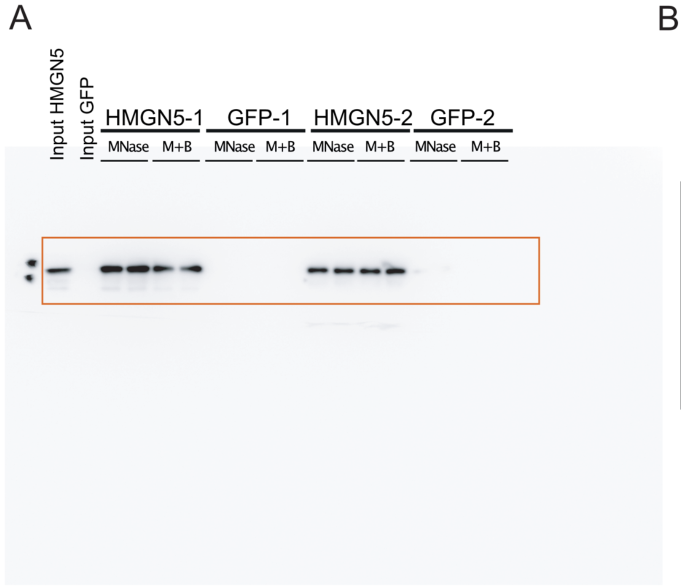


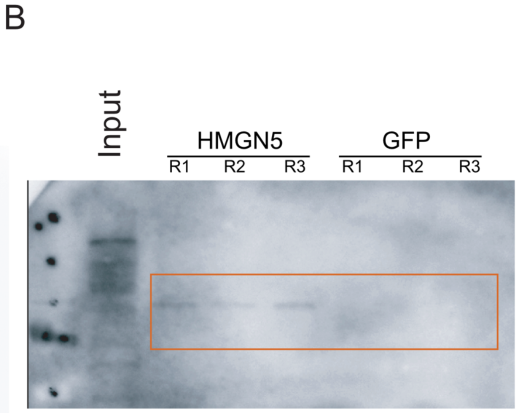


**Supplementary Figure 9.** Uncropped western blots of pictures presented in Figure 7c (A) and Figure 7e (B).

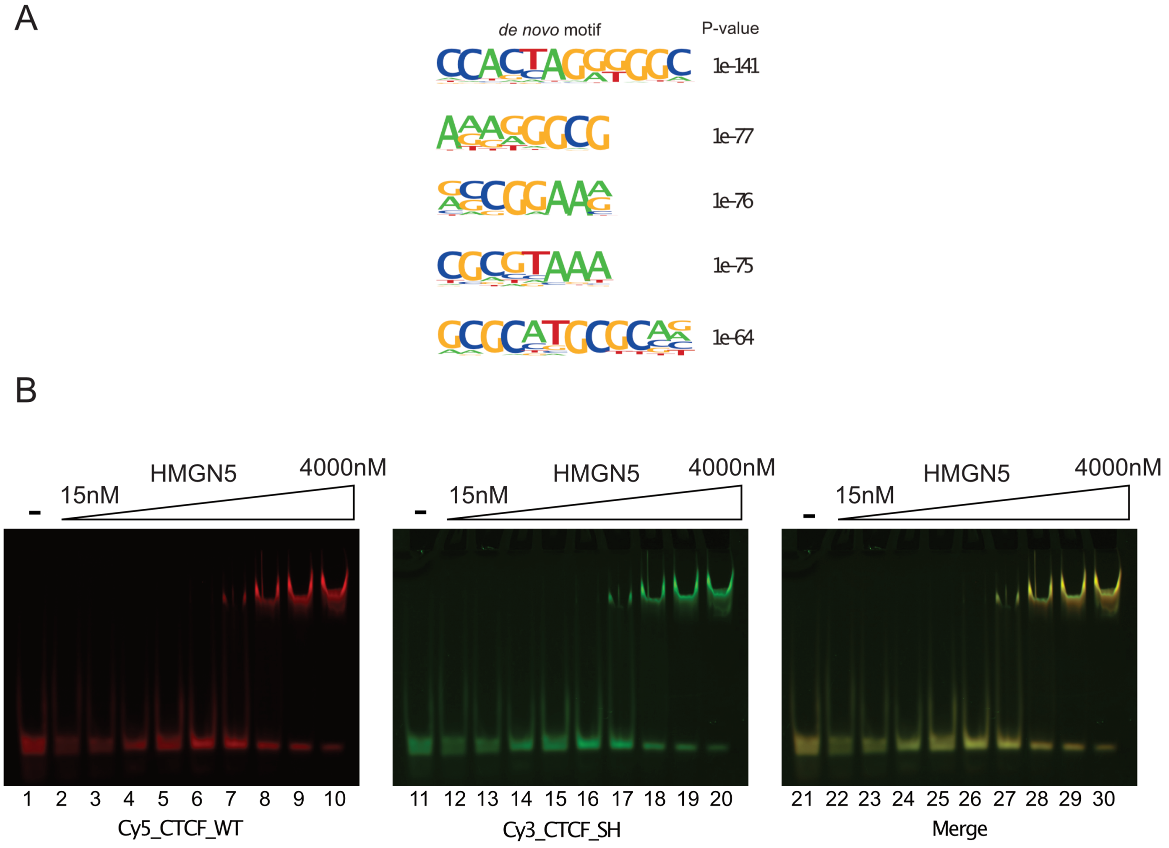


**Supplementary Figure 10.** (A) *De novo* motif analysis using the HMGN5 ChIP-Seq peaks. Top five most enriched motifs in the HMGN5 ChIP-seq dataset are ranked according to the p-value. The motifs were determined with the software Homer assuming a peak size of 1000bp. (B) Interaction of HMGN5 with the double stranded CTCF wild type motif (Cy5_CTCF_WT) and the mutated motif (Cy3_CTCF_SH). The Cy5 and Cy3 labeled DNA molecules were mixed at equimolar ratios and subsequently incubated with increasing HMGN5 concentrations as indicated. The individual binding reactions are visualized in lanes 1-10 (CTCF wild type motif) and lanes 11-20 (CTCF mutated motif). A merge of both gels is shown in the right panel.

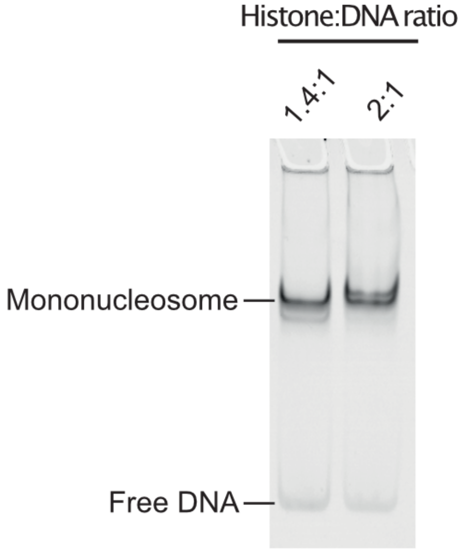


**Supplementary Figure 11.** *In vitro* assembly of mono-nucleosomes. Mono-nucleosomes were reconstituted using a Cy3-labeled PCR fragment containing the 601 sequence, and histones purified from chicken blood. The assembly was performed by salt dialysis method using a histones:DNA ratio of 1.4:1 and 2:1. Reconstituted nucleosomes were analyzed by native gel electrophoresis (6% PAA, 0.4x TBE Buffer) and documented with a fluorescence imager (Thyphoon 9500).

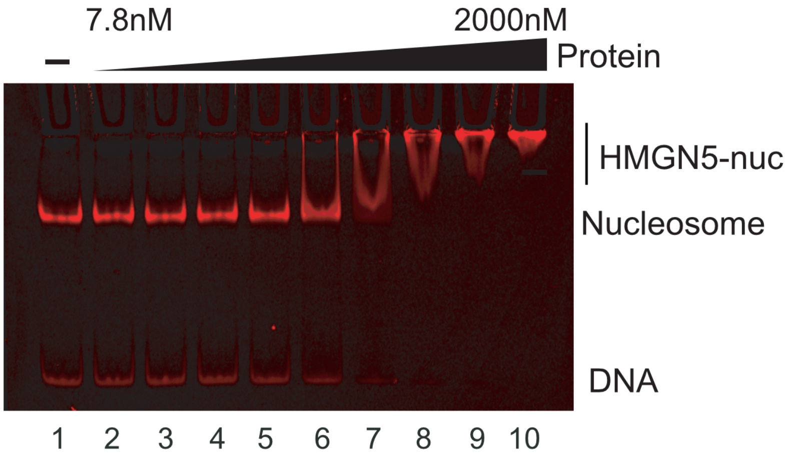


**Supplementary Figure 12**. Analysis of HMGN5 mono-nucleosome interaction. Reconstituted nucleosomes (Cy3-labeled) were used at a concentration of 50ng per reaction (lanes 1 to 10). HMGN5 Protein was added in increasing concentrations, starting at 7.8nM and increasing 1.5 fold up to 2000nM, as indicated (lanes 2 to 10). Reactions were analyzed on 6% PAA gels, electrophoresed in 0.4x TBE buffer and documented using fluorescence imager Typhoon FLA 9500 (GE Healthcare).
